# Local and distal signals drive cell-type-specific leaf responses during shade avoidance

**DOI:** 10.64898/2026.10.11.757655

**Authors:** Lisa Oskam, Leonard Jo, Carolin Grones, Muthanna Biddanda-Devaiah, Nicole M. Nieuwendijk, Ronald Pierik

## Abstract

Plants reposition and grow their leaves towards optimal light conditions in a canopy. To achieve both horizontal and vertical displacement, plants need to simultaneously elongate their cells unidirectionally and maintain an elongation rate differential between the abaxial and adaxial side of the organ to bend. Exposure to Far-Red (FR) light enrichment can induce both responses, depending on where FR is sensed. Here we studied how plants maintain both growth responses simultaneously within the same organ. To deduce the regulatory mechanisms governing the simultaneous regulation of differential and unidirectional elongation, we performed both organ specific and single cell RNA sequencing experiments on plants exposed to local or whole plant FR enrichment. Single cell transcriptomics resolved that FR light signalling occurs through all cell types, but core components of the shade avoidance pathway are expressed cell type-specifically and require signalling across tissues to control growth plasticity. The upstream photoreceptor targets, including auxin biosynthesis-associated genes, in the leaf lamina are expressed in the epidermis, whilst auxin response gene expression in the petiole occurs also in the deeper tissues layers. We thus identified the remotely triggered differential growth transcriptome acting in the abaxial cortex and show that this is complemented by locally induced second transcriptome signature that is associated with unidirectional petiole elongation. The latter is induced across all cell types. We propose that this spatial distinction of transcriptional regulation of gene clusters could represent how differential and unidirectional growth can occur simultaneously in the same organ under FR light enrichment.

## Introduction

Plants compete for resources when growing in dense populations such as in agricultural fields. Competition for sunlight occurs through shade avoidance responses, where plants reposition their foliage away from shade that is cast by neighbouring competitors to ensure optimal light capture (Casal and Fankhauser 2023). Rosette plants such as *Arabidopsis thaliana* respond by both elongating their petioles and moving their leaves upwards (Fernández-Milmanda and Ballaré 2021), whereas vegetative stem-forming plants also strongly elongate their internodes (Li et al. 2024). True shade consists of both a reduced light intensity over the full light spectrum, measured in Photosynthetically Active Radiation (PAR), and a low Red to Far Red light ratio (R:FR) (Ballaré et al. 1990). Neighbouring plants enrich the spectrum with FR, due to leaf FR reflection, causing a lowering of the R:FR ratio even before a reduction in PAR is perceived (Huber et al. 2021). FR enrichment functions as an early indicator of encroaching competition. Light distribution in canopies is highly dynamic and heterogeneous and so is the spectral composition (Durand and Robson 2023). Plants in competition thus rarely experience homogeneous shade cues, and it was shown previously that depending on where on the plant body FR enrichment occurs, different aspects of the shade avoidance syndrome are elicited. FR enrichment on the leaf lamina tip is necessary and sufficient to elicit upward movement of the petiole (Michaud et al. 2017, Pantazopoulou et al. 2017), also known as hyponasty, whereas local FR enrichment on the petiole elicits petiole elongation but not hyponasty (Pantazopoulou et al. 2017). Whole plant exposure to FR enrichment induces both unidirectional petiole elongation and hyponasty simultaneously (Michaud et al. 2017, Pantazopoulou et al. 2017, Oskam et al. 2024). Hyponasty emerges from differential growth rates between the adaxial and abaxial side of the petiole, leading to slight upward bending of the petiole base, thus moving the leaf to a more upward orientation (Polko et al. 2012, Küpers et al. 2023). Organ bending often involves cell elongation on the side opposite of the bending direction (Jonsson et al. 2023), as is happening during hyponasty. However, during whole plant FR enrichment, both the abaxial and the adaxial side of the petiole need to elongate to drive unidirectional petiole elongation that co-occurs with hyponasty (Pantazopoulou et al. 2017, Oskam et al. 2024), while also maintaining differential elongation between the abaxial and adaxial side.

The molecular regulation of the hyponastic response has been partially elucidated. Low R:FR inhibits phytochrome B (phyB) activity, releasing the inhibition of PHYTOCHROME INTERACTING FACTORS (PIFs) (Cheng et al. 2021b). PIF7 is the main PIF required for local FR-induced hyponasty, together with PIF4 and PIF5 (Pantazopoulou et al. 2017, Küpers et al. 2023). PIFs are basic Helix-Loop-Helix (bHLH) transcription factors, and enhance *YUCCA (YUC)* expression, inducing auxin biosynthesis (Sellaro et al. 2017). Auxin biosynthesis enzymes YUC2, YUC5, YUC8 and YUC9 are all involved in the hyponastic response, with *YUC8* and *YUC9* showing strong induction in the lamina upon FR on the leaf tip (Küpers et al. 2023). Auxin is transported in a PIN3- , PIN4- and PIN7-dependent manner towards the abaxial side of the petiole base, where it induces differential elongation (Michaud et al., 2017; Pantazopoulou et al., 2017; Küpers et al., 2023). Other hormones such as gibberellic acid (GA) and abscisic acid (ABA) play a role either in finetuning of the elongation response or in general leaf angle setpoint (Küpers et al. 2023, Michaud et al. 2023). Brassinosteriods and gibberellins imping on petiole elongation through the putative BRASSINAZOLE RESISTANT 1-AUXIN RESPONSE FACTOR 6-PHYTOCHROME INTERACTING FACTOR 4/DELLA (BAP/D) transcription factor module (Oh et al. 2014). Despite the knowledge on this basic network, it is currently unknown which cell layer(s) regulate the co-occurring phenotypic FR responses. Regulation of organ elongation has been attributed to the epidermis in hypocotyls (Savaldi-Goldstein et al. 2007, Procko et al. 2016, Kelly-Bellow et al. 2023). However, regulation of leaf angle setpoints regardless of supplemental FR through ABA was described in the mesophyll layer (Michaud et al. 2023). Autotropism, or the straightening of a previously bent organ, on the other hand is proposed to be controlled through xylem fiber cells (Okamoto et al. 2015). The initial perception of a change in the R:FR ratio could be in any cell layer of the lamina, as phyB is expressed ubiquitously throughout the plant (Goosey et al. 1997), but tissue layer specific regulation has been described for phyB in relation to flowering, where mesophyl-expressed phyB suppressed FLOWERING LOCUS T expression in the vascular bundle (Endo et al. 2005).

Although previous research has uncovered major parts of the regulation of the hyponastic response at an organ specific level, it is currently unknown which cell layers regulate which steps of the regulatory cascade in both the lamina and the petiole. Understanding this task division across tissues may help resolve how the leaf balances unidirectional and differential elongation when not only the leaf tip experiences supplemental FR, but the petiole as well. To study the spatial specificity of leaf shade avoidance response, we performed single cell and bulk RNA sequencing experiments on dissected leaves. The scRNAseq experiment showed that although *phyB* was expressed and active throughout all cell types, *PIF7*, *YUC8*, and *YUC9* expression and regulation by FR in the lamina was mainly restricted to the epidermis. The resulting light-induced auxin that is translocated to the petiole elicits transcriptional responses in the deeper abaxial tissues of the petiole as well as in the epidermis. These findings highlight across-tissue layer communication for hyponasty. Complementing these patterns are transcripts whose expression is induced on both abaxial and adaxial sides and combing the different sequencing experiments we identify a petiole elongation gene set that acts in concert with the hyponasty gene set. The functional and tissue-specific separation of the hyponasty and elongation gene sets likely underpins how a leaf can control two adaptive behaviors to an environmental cue simultaneously.

## Results

### FR signalling is activated in all cell layers, whereas shade avoidance targets respond cell type-specifically

To investigate how plants process neighbour proximity cues to produce growth adjustments, we performed a single cell RNA sequencing (scRNAseq) experiments on dissected leaves, separating lamina from petiole. Hyponasty is induced by signalling from the lamina tip (Pantazopoulou et al. 2017)(Supplemental Figure 1A), whereas petiole elongation can be induced upon signalling of local FR enrichment in the petiole itself (Pantazopoulou et al., 2017). Homogeneous whole plant FR enrichment elicits both responses simultaneously (Oskam et al. 2024)(Supplemental Figure 1A), and is used here to resolve whether these co-occurring responses involve differential patterns and cell types.

Laminas and petioles were harvested, protoplasted and processed separately after either three hours of FRw (FR-enriched white light on the whole plant) treatment, or white light (WL) control treatment (Figure 1A). After the integration of datasets from the WL and FRw enriched samples, cells were clustered to identify distinct cell types in the respective tissues (Figure 1B, C). Cells clustered for cell type rather than treatment for both lamina and petiole cells (Figure 1B, C). Lamina and petiole derived cells clustered into 22 and 25 groups, and clusters were attributed to 14 and 12 different cell types respectively using the expression patterns of a previously established cell-type marker gene set (Supplemental Figure 1B, C; (Tenorio Berrío et al. 2022). For the final cell type annotation, marker gene expression as well as cluster information was combined into cell type subdivisions that could potentially capture cell states (Supplemental Figure 1D). To identify genes that were responding to the FRw enrichment treatment, we compared gene expression levels in cells in WL to FRw per individual cell cluster for both tissues (Figure 1D, E, Supplemental Table 1,2). The numbers of up or down differently expressed genes (DEGs) were comparable between petiole and lamina clusters, but highly variable between the different cell types.

**Figure 1.**
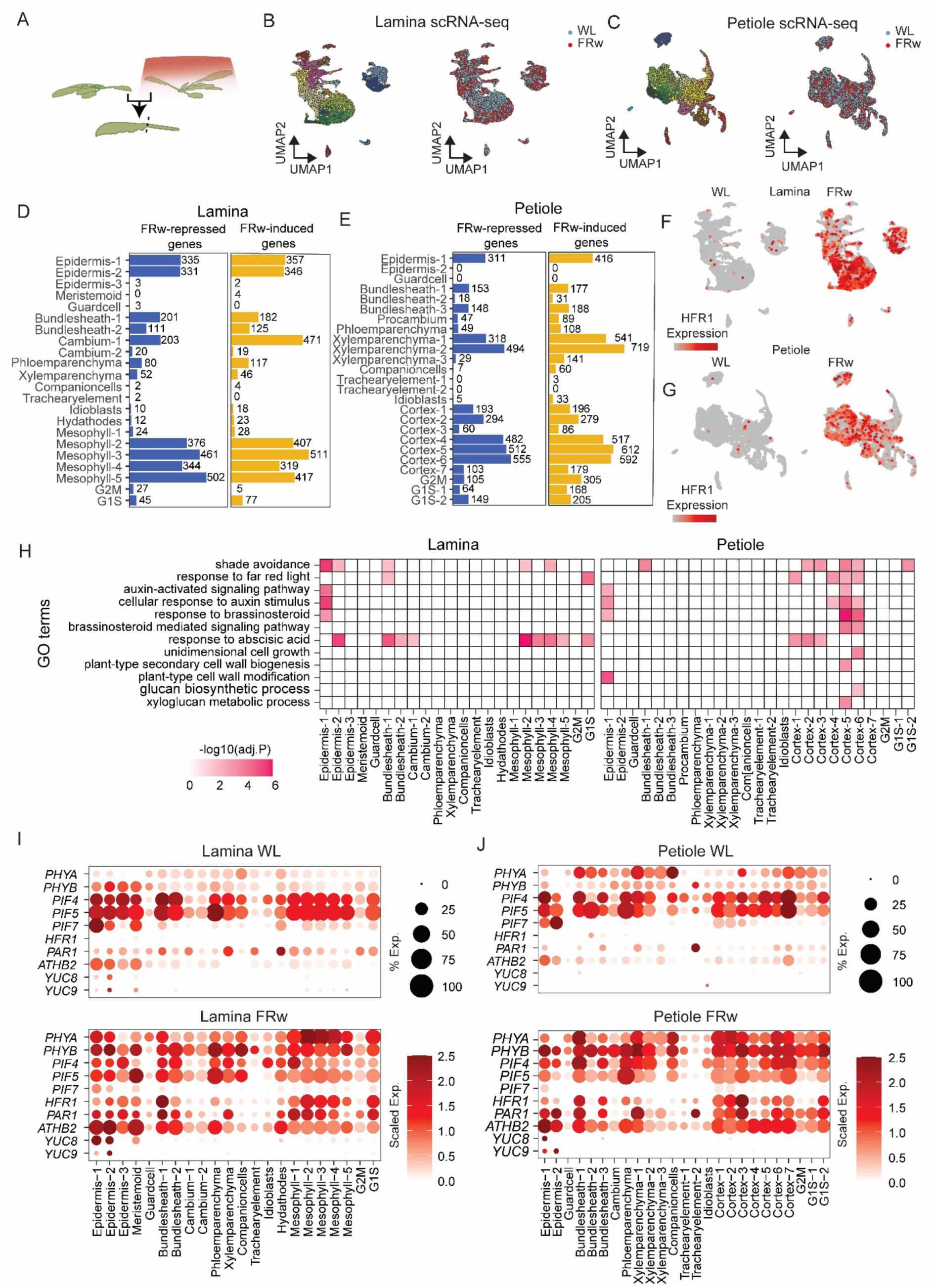
Tissue specific expression of whole plant far-red (FRw) enrichment-induced genes. (A) Experimental set-up for single cell RNA sequencing (scRNAseq) of laminas and petioles of plants exposed to far-red enrichment (FRw) or white light control (WL) conditions. (B) Uniform Manifold Approximation and Projection (UMAP) for both the different clusters and WL or FRw treated tissue of the lamina and (C) the petiole. (D) Differentially expressed genes (DEGs) for the different cell type clusters, either FRw-repressed in blue, or FRw-enhanced in yellow in the lamina, and (E) in the petiole. (F) Expression of FR-induced marker gene HFR1 in WL and FRw represented in a UMAP plot of the lamina, and (G) the petiole. (I) Both the relative expression and percentage of cells expressing the gene in question per cluster, of shade avoidance related genes in the lamina in WL and FRw, and (J) in the petiole.

The established FR enrichment signalling marker *HFR1* was FR-induced across most cell type clusters in both the lamina and the petiole (Figure 1F, G). A gene ontology (GO) enrichment analysis of cell type up regulated DEGs highlighted shade avoidance regulation in epidermis, bundle sheath and mesophyll/cortex of petiole and lamina, whereas response to auxin in the lamina is restricted to the epidermis, but especially strong in the cortex of the petiole (Figure 1H). Similarly, other canonical FR enrichment signalling indicator genes such as *KDR*, *PIL1*, *PAR1*, *PAR2*, and photoreceptor *PHYB* showed enhanced expression under FRw enrichment throughout the majority of cell types (Figure 1J, K). *PIFs* transcript levels are repressed in FRw enrichment, consistent with their established negative feedback loop of transcriptional self-repression (Leivar and Monte 2014). In both the lamina and the petiole, *PIF4* and *PIF5* are expressed throughout the different cell types (Figure 1J, K). *PIF7* is specifically associated with hyponasty and displays a less generic expression pattern, with strong bias towards the epidermis, and a mild expression in mesophyll and cortex clusters (Figure 1J, K). Expression of direct targets of PIF7, *YUC8* and *YUC9* is even more locally restricted to the epidermis clusters (Figure 1J, K).

Together, the regulation of the shade markers and GO enrichment indicate that FR enrichment is likely perceived throughout the entire leaf and several direct downstream targets of phyB are activated in all cell types. However, auxin regulation appears to be more specified towards specific tissues in the lamina and petiole.

### Leaf sections show distinct gene expression response to local and systemic FR enrichment

Although the scRNAseq offers great potential for studying gene expression in cell types, deeper insights between the abaxial and adaxial petiole sides were required to interpret hyponasty. To this end, we performed a bulk RNAseq experiment in which we physically separated the petiole into an abaxial and adaxial side during sampling, in addition to sampling the lamina tip. We included an additional treatment that elicits only hyponasty, not elongation: spotlight FR enrichment of the lamina tip (FRt) (Figure 2A, Supplemental Table 3; (Oskam et al. 2024). The up and down regulated FRw related DEGs showed a high overlap with the cell cluster specific DEGs from the lamina and petiole (Supplemental Figure 1E, F). Focusing on upregulated DEGs of the bulk RNAseq, we identified 255 DEGs overlapping between FRt and FRw in the lamina (Figure 2A). An even larger subset of 412 genes was specifically upregulated in supplemental FRw and thus not in FRt (Figure 2A), and a comparable number of downregulated genes was specific to FRw (376) (Supplemental Figure 2A). A GO enrichment analysis of the DEGs in the three parts of the Venn diagram showed that the main GO categories associated with shade avoidance, FR light and auxin responses were found to be regulated by both local FRt and FRw enrichment in the leaf tip (Supplemental Figure 2B).

**Figure 2.**
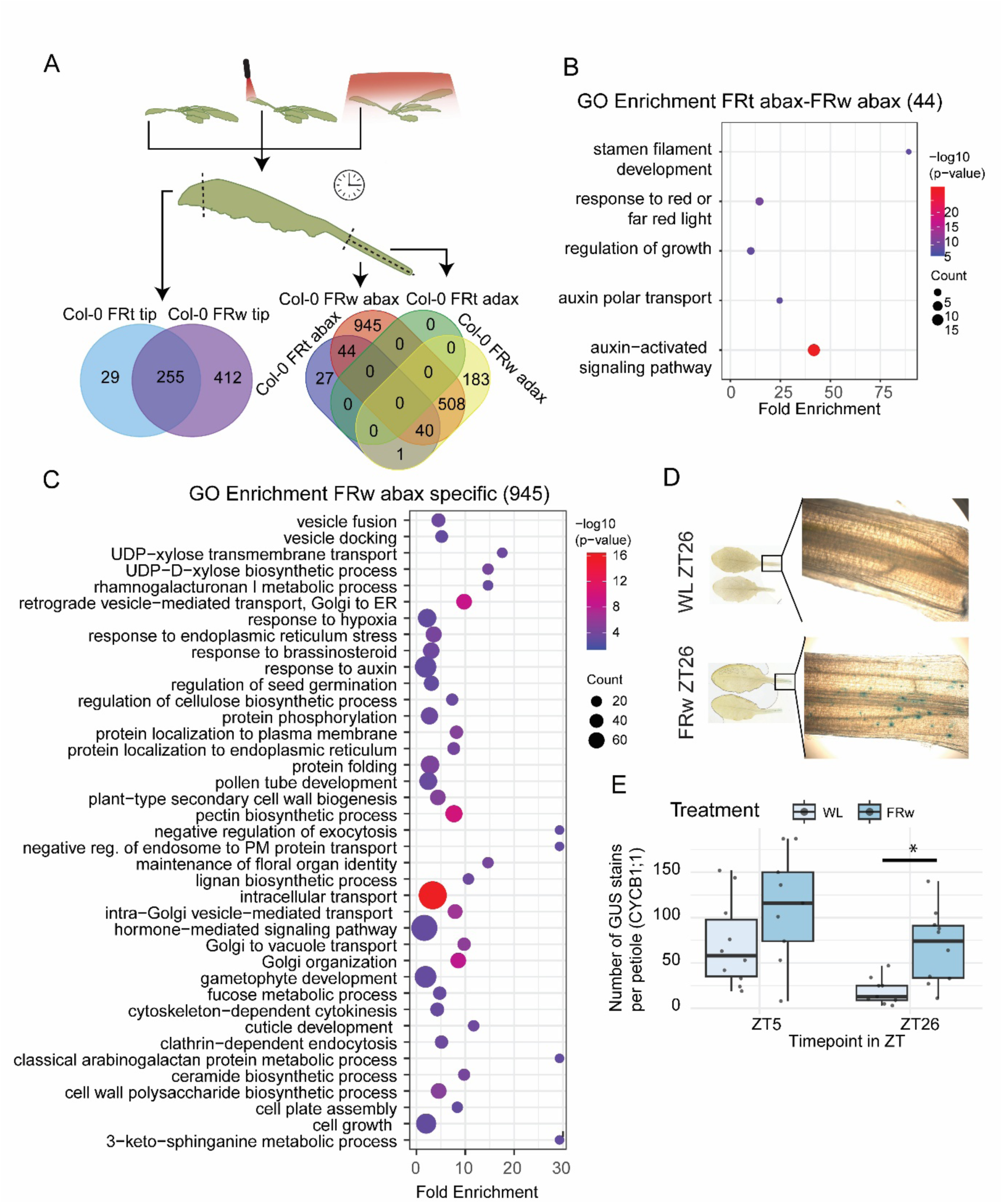
Local FRt and whole plant FRw enrichment induce a common shade avoidance response, while revealing a distinct program for abaxial cell elongation. (A) RNA sequencing design of Col-0 after a 3-hour treatment of either WL, supplemental FRt or supplemental FRw. The leaf tip, and two thirds of both adaxial and abaxial petiole section closest to the rosette centre were harvested separately. Venn diagram of upregulated DEGs of the leaf tip and the petiole. (B). Enriched GO categories of DEGs overlapping between FRt and FRw enrichment in the abaxial petiole. Three biological replicates per treatment, tissue and genotype combination were analysed. (C) GO categories derived from the subset of 508 DEGs that were specifically upregulated in supplemental FRw in the abaxial petiole. (D) Scans and stereoscope image (10x) of petioles after FRw or WL treatment at ZT26, treatment start time at ZT2. (E) Number of GUS stains per petiole image per treatment and ZT, t-test was done per ZT timepoint, 10 replicates per treatment and timepoint.

In the petiole, there were 112 DEGs on the abaxial side under FRt treatment, as compared to 1537 DEGs in the same tissue under FRw treatment, indicating how signalling from a remote tissue has a far more limited effect than local exposure and signalling (Figure 2A). However, the contrast between ab- and adaxial petiole gene expression induced by FRt treatment of the lamina was stark (0 DEGs adaxial versus 112 DEGs abaxial; Figure 2A). The subset of 44 DEGs that overlap between FRt and FRw treatment specific to the abaxial petiole was enriched in several auxin and FR-related GO categories (Figure 2B). We propose this gene set as the core hyponasty genes, firstly, because their expression is regulated across both FR-enrichment treatments that induce hyponasty. Secondly, their expression is spatially restricted to the abaxial side, where growth promotion has to occur specifically for differential growth (see Supplemental Figure 2C for expression levels across the different treatments). This core hyponasty gene set is supplemented by 40 DEGs that are associated with elongation and whose expression is FR-induced in any elongating tissue: the abaxial petiole in FRt, and both the abaxial and adaxial petiole during FRw, where the combination of abaxial and adaxial elongation contributes to unidirectional petiole elongation (see Supplemental Figure 2D for expression levels across the different treatments).

The subset of 945 DEGs that were specifically upregulated in supplemental FRw in the abaxial petiole were enriched in a plethora of GO categories related to intracellular trafficking pathways, but also categories such as those related to cell wall modifications, cytokinesis, cell growth and cell differentiation (Figure 2C). Furthermore, the auxin biosynthesis double *yuc8 yuc9* and quadruple *yuc2 yuc5 yuc8 yuc9* mutants had reduced or absent hyponasty upon FR enrichment but they did maintain the elongation response (Supplemental Figure 3A-D). These observations indicate that auxin alone may not control petiole elongation under FRw enrichment. Based on the GO enrichments for cytokinesis we used the CYCLINB1;1 (CYCB1;1-GUS) (Eloy et al. 2011) reporter for cell cycle activation and found that it remained more active in response to FRw in the petiole (Figure 2D, E). At ZT5, the time corresponding to the time of the RNAseq, both WL and FRw treated petioles displayed GUS activity (Figure 2E). However, at ZT26, the FRw enriched petioles showed a higher CYCB1;1-GUS activity around the vasculature and cortex regions.

### Defining abaxial and adaxial cell clusters identifies tissue-specific patterns of shade avoidance genes

Next, we used the abaxial and adaxial petiole samples from the manually dissected bulk RNAseq to identify transcripts that are strongly enriched in either of the two fragments, aiming to use these to further refine the cell clusters of the scRNAseq experiments. We selected the top 20 genes with the strongest abaxial or adaxial enrichment (Figure 3A) in WL to score each cell in the petiole scRNAseq dataset into abaxial, adaxial, or undefined (Figure 3B). The dashed line in Figure 3C indicates where the abaxial section was manually split from the adaxial section during sample harvest for bulk RNAseq. High confidence scoring was observed for the seven cortex clusters (Figure 3D). Cortex 1-3 had a very strong adaxial signature, whereas cortex 5-7 had a strong abaxial signature. Cortex 4 displayed an equal distribution of adaxial and abaxial cells. This demonstrates that some cell-types retain their positional identity, which is reflected in their transcriptional profiles. Vascular bundle tissues were also identified predominantly as abaxial, but we will treat these clusters separately as vasculature as the assignment to abaxial was likely a result of the broader expression pattern of the genes chosen as abaxial versus adaxial markers (Supplemental Figure 3E). Expression of shade marker *PIL1* occurred throughout the cortex subclusters (Figure 3E), while expression of auxin-response markers like *SAUR19*-*24* was restricted to abaxial cortex subclusters (Figure 3F, G).

**Figure 3.**
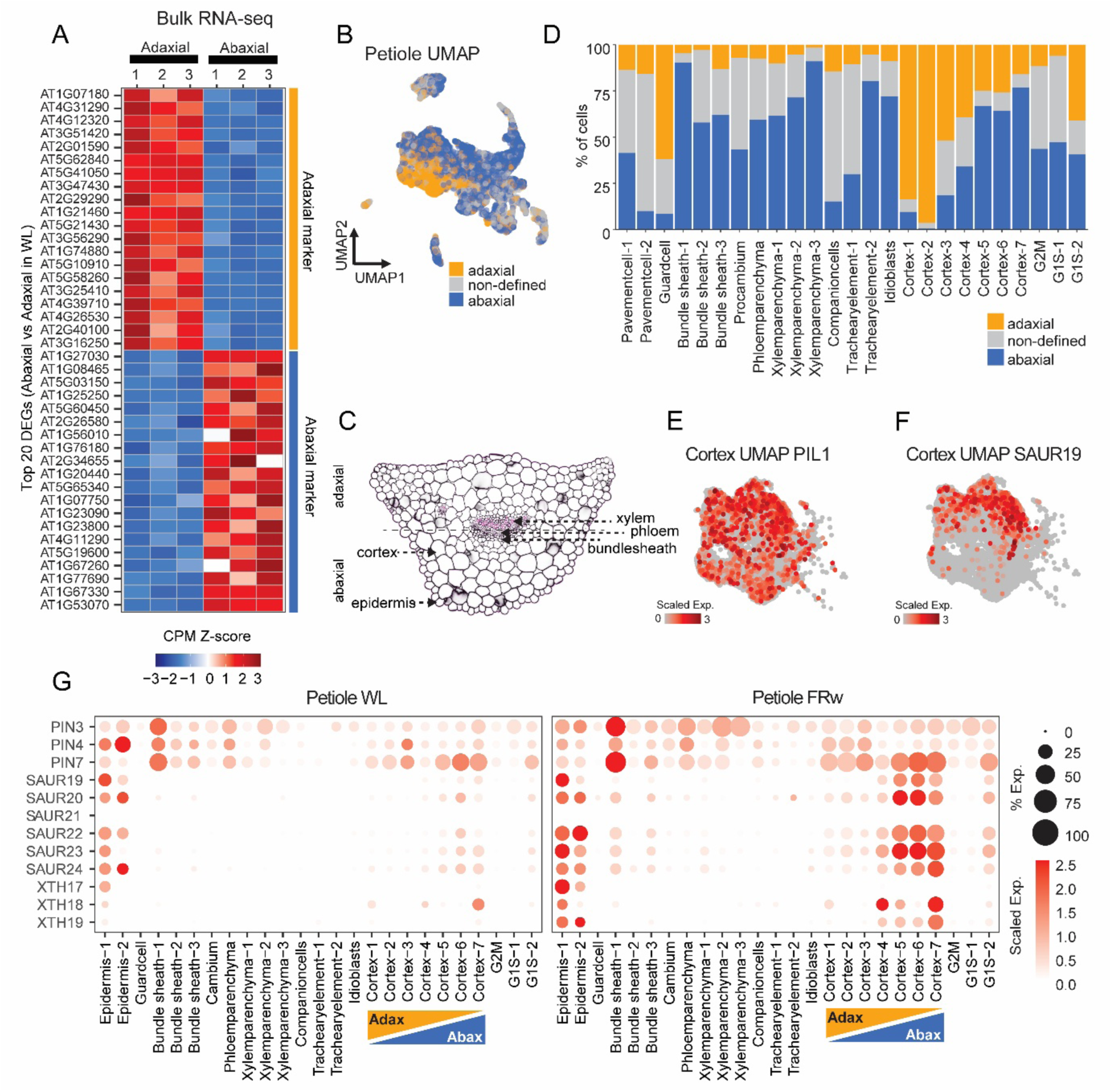
Petiole cortex clusters can be assigned to abaxial or adaxial positioning in the petiole based on manually dissected and RNA sequenced tissue. (A) Z-score of the Counts Per Million (CPM) of the 20 most upregulated genes in either abaxial or adaxial petiole tissue from the three WL treated samples. (B) Petiole UMAP of the different cells assigned to either abaxial, adaxial or non-defined based on the marker shown in A. (C) Confocal cross section of the petiole at around 500 µM from the petiole base. Pink colouring indicates xylem tissue through Basic Fuchsin (0.2%) staining. (E) UMAP of cortex clusters 1-7 of PIL1 and (F) SAUR19 in FRw enrichment. (G) Expression and percentage of cells expressing genes related to auxin transport and response, and cell wall modifying related genes in WL and FRw enrichment.

With the further distinction of cortex clusters into the abaxial or adaxial side of the petiole, we continued to explore the gene expression patterns throughout the different tissue type clusters. We first looked deeper into expression patterns of the shade avoidance pathway through sets of previously established genes (Oh et al. 2014, Kohnen et al. 2016, Michaud et al. 2017, Pantazopoulou et al. 2017, Küpers et al. 2023, Michaud et al. 2023)(Supplemental Figure 4 – 6). The majority of genes related to early signalling and transcriptional control upon FR enrichment were expressed in all cell types in both petiole and lamina (Supplemental Figure 4A-D). This was already the case in control WL, and holds for the induction of expression by FR enrichment, which overall induced higher gene expression in the petiole compared to the lamina. Genes related to the hormones auxin, gibberellin and abscisic acid displayed a more localized expression pattern in WL (Supplemental Figure 5,6). FRw enrichment promoted gene expression in the petiole compared to the lamina for IAA and specific GA-related genes, but less obviously for ABA-related transcripts. These variations in tissue localization of FRw-induced transcripts hint at potentially diverging roles for epidermis, cortex and vasculature bundle for hormone-related events in shade avoidance.

### Coordination of unidirectional and differential petiole elongation is associated with spatially distinct transcriptional programs

With the established petiole abaxial and adaxial cortex distinction we took the core hyponasty set of 44 genes (Figure 2A) and visualized their expression in the scRNAseq experiment in the petiole (Figure 4A). Although regulation occurred across tissue layers, a strong bias of *SAUR* auxin response genes was found in the abaxial cortex. (Figure 4A). The second subset of 40 genes in the general elongation gene set (Figure 2A), showed a very different pattern with an induction by FR enrichment in epidermis, vasculature bundle and cortex clusters (Figure 4B). Despite this more generic induction across all cell types, the induction was still substantially stronger in the abaxial compared to adaxial cortex clusters.

**Figure 4.**
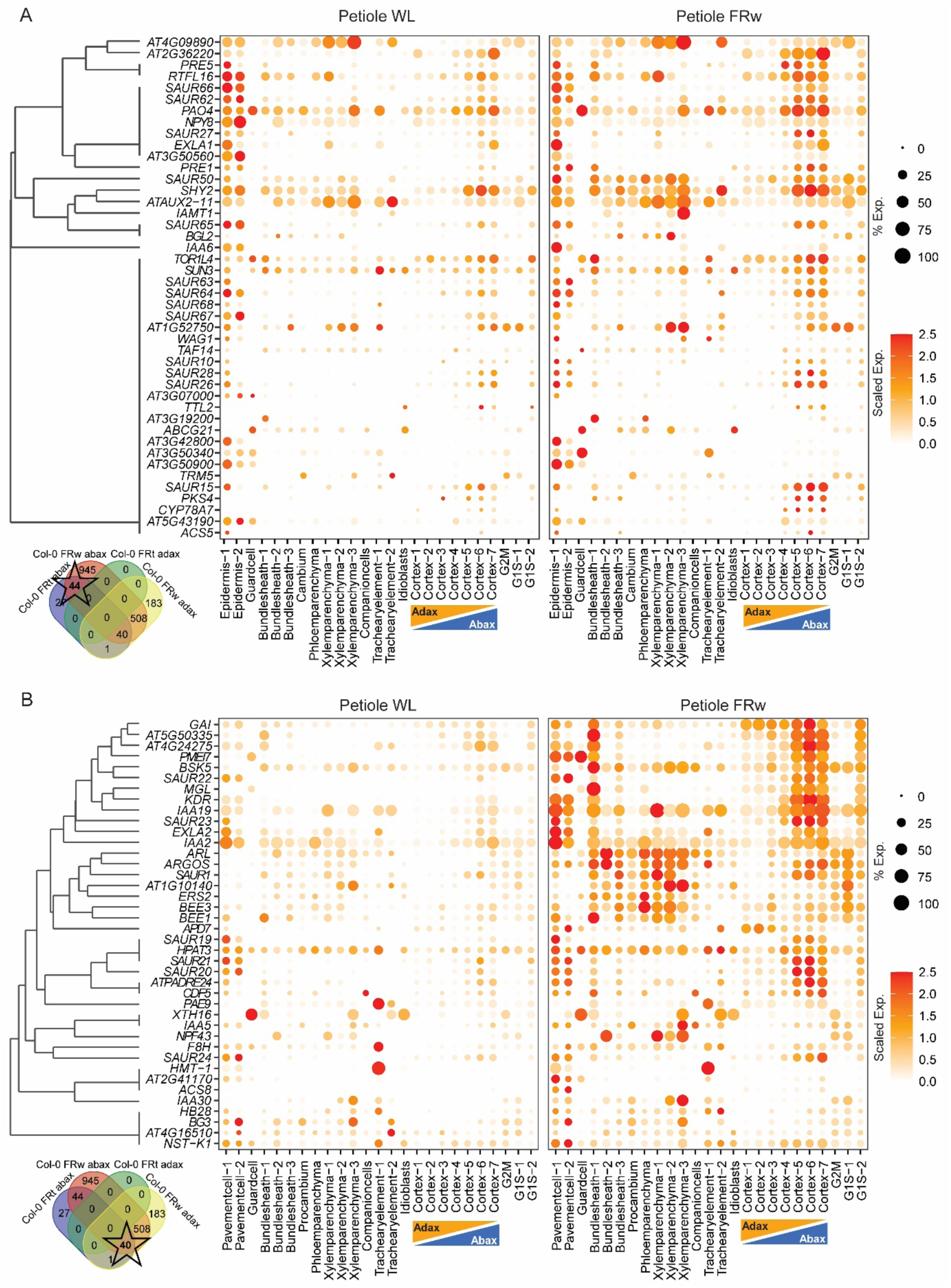
Hyponasty-associated transcriptional reprogramming is enhanced in the abaxial cortex layer while FR-induced elongation-related transcriptional reprogramming is occurs across cell types . (A) Average gene expression level and relative expression per cluster of DEGs from FRt and FRw enrichment bulk RNAseq experiment plotted in the FRw enrichment scRNAseq for the petiole of 44 hyponasty core set DEGs and (B) of 40 general elongation core DEGs. Colour scale represents average level of expression per gene, set from 0.0 to 2.5 max. Size of the dot represents the percentage of cells in a cluster expressing the gene in question. Gene clustering was based on their cell-type-specific differential expression pattern, where the differential expression was encoded as a binary variable (non DEG = 0, DEG = 1). Genes were then grouped by hierarchical clustering using the binary distance (Jaccard).

To further resolve the petiole elongation response, that requires elongation on both the abaxial and adaxial side, we performed another independent bulk RNAseq experiment in which we compared control WL, whole plant FR (creating hyponasty and unidirectional petiole elongation) and spotlight FR illumination of only the petiole (creating unidirectional petiole elongation but not hyponasty) (Figure 5A). Since we observed that the petiole elongation response to local FR enrichment depends strictly on PIF4 and PIF5, we performed this experiment in both wildtype Col-0 and the *pif4 pif5* double mutant (Figure 5B). Consistent with a stronger petiole elongation phenotype, FRw treatment let to a much larger number of DEGs in the petiole than did FRp, but of the 180 FRp-upregulated DEGs, 155 were shared with FRw-upregulated genes (Figure 5A,B). We then took these petiole DEGs that are shared between FRp and FRw, and compared these with the FRw DEGs in the previous RNAseq experiment where we had split the petiole longitudinally in abaxial and adaxial fragments (Figure 5C). Specifically, we overlayed the 155 petiole elongation-related DEGs with 508 DEGs for genes that were FRw-induced in both abaxial and adaxial fragments and not in the abaxial petiole during FRt treatment. We continued with the overlap since these genes should be most strictly associated with unidirectional petiole elongation. After these steps, 70 DEGs were left, and we term these the core FR-induced petiole elongation genes (see Supplemental Figure 7 for expression levels across the different treatments), complementary to the previously identified core FR-induced hyponasty genes and the general elongation genes (Figure 5C, D). Additionally, we compared this set with the list of DEGs in FRp in the *pif4 pif5* double mutant (Supplemental Table 4). Out of the 70 core FR-induced petiole elongation genes, 60 were not differentially expressed in the *pif4 pif5* double mutant, suggesting that these genes are dependent of PIF4 or PIF5 in response to far-red light in the petiole (Figure 5C, D). Figure 5D shows tissue-specific expression of the unidirectional petiole elongation core set under WL and FRw enrichment. In contrast to the hyponasty and general elongation gene sets, the unidirectional elongation core set was induced across nearly all cell clusters and lacked the bias towards abaxial cortex expression. This was especially pronounced in the epidermis, where the unidirectional petiole elongation core set showed strong expression, while the hyponasty core set was not differentially expressed. This suggests that the FR-induced petiole elongation occurs across cell types, while FR-associated hyponasty responses are triggered especially in the abaxial cortex. This could imply that the cortex plays a major role in defining differential growth, thus specializing regulation of the two phenotypes across tissues. In addition, the expression of the general elongation core set (Figure 4B) also occurred in the epidermis, vasculature, as well as the abaxial cortex, in FRw enrichment. This suggests that the abaxial cortex shows a strong transcriptional signature associated with differential growth, which is supplemented with coordinated regulation across multiple cell types.

**Figure 5.**
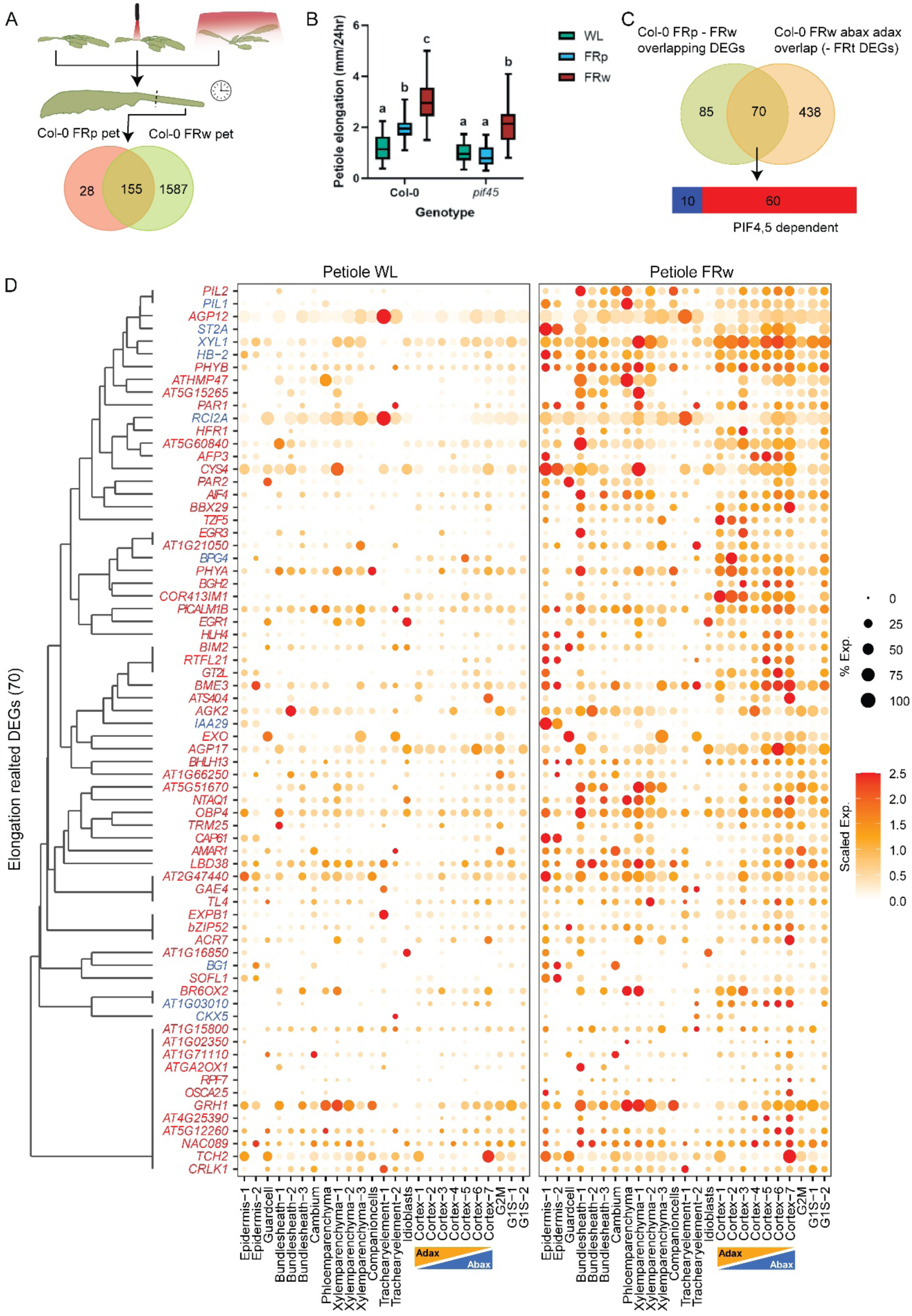
Petiole elongation is associated with gene expression patterns across most cell types. (A) Bulk RNAseq and corresponding DEGs for COL-0 with WL control, local FR petiole (FRp) and whole plant FR (FRw) enrichment, petioles were harvested after 3 hours of treatment. (B) Phenotypic responses of Col-0 and pif4 pif5 double mutant to WL, FRp and FRw enrichment. (C) Filtering of genes using 508 DEGs derived from the bulk RNAseq on dissected tissues (Fig. 2) that were specific for FRw treatment in abaxial and adaxial tissue. These genes were further filtered based on their expression in the pif4 pif5 double mutant bulk RNAseq data. (D) Average gene expression and percentage of cells expressing a gene of the unidirectional petiole elongation core set per cluster in the petiole for both WL and FRw enrichment. Gene clustering was based on their cell-type-specific differential expression pattern, where the differential expression was encoded as a binary variable (non DEG = 0, DEG = 1). Genes were then grouped by hierarchical clustering using the binary distance (Jaccard). Colour scale represents average level of expression per gene, set from 0.0 to 2.5 max. Size of the dot represents the percentage of cells in a cluster expressing the gene in question.

### Enhanced growth regulation patterns associated with combined shade avoidance traits

While both the core hyponasty and core petiole elongation gene sets define the genes associated with either of the traits, the goal is to identify how these phenotypic response can be expressed simultaneously. To this end, we took all genes that were DEGs and overlapping between FRw abaxial and adaxial and no other tissues, and the DEGs that were overlapping between FRw ab-ad and with FRt abaxial (Figure 2A). This set of 548 genes was filtered by removing the DEGs that were overlapping with DEGs that were present in FRt and FRw treated lamina’s (Figure 6A), focusing the gene set on response regulation rather than light signalling itself. Since FRp only induced mild elongation compared to FRw enrichment, we postulate that FRw induces additional patterns associated with the additional elongation. We therefore, also filtered against genes that were already induced by FRp treatment, leaving DEGs that are explicitly regulated additionally during whole plant FR enrichment where petiole elongation and hyponasty co-occur, and are likely associated with control of these phenotypes (Figure 6B). This gene set describes patterns that occur in the petiole during whole plant FR enrichment that are specific to this whole-plant treatment, thus excluding the core petiole elongation and hyponasty genes that are regulated also by the highly localized petiole and leaf tip spotlight FR treatments. Two broad enrichments spread over multiple GO terms that stood out were additional hormonal regulation, particularly brassinosteroids (BR) and gibberellin (GA), and growth regulation (Figure 6B). Growth regulation related categories contained DEGs that were mostly upregulated in the abaxial cortex, vasculature and epidermis (Figure 6C). GA and BR related DEGs were upregulated across all tissues, including adaxial cortex, but their expression levels were already relatively high in WL control (Figure 6C). As expected from the selection criteria, transcript levels of the selected genes were not much affected by light in the lamina, or in other treatments (Supplemental Figure 8). Previous studies have indicated the epidermis as the main site of control for elongation regulation including by brassinosteroids (Savaldi-Goldstein et al. 2007, Hacham et al. 2011), although this idea has also been challenged (Blanco-Touriñán et al. 2024). The observed expression profiles indicate a very broad BR and GA related transcriptional response across all tissue layers.

**Figure 6.**
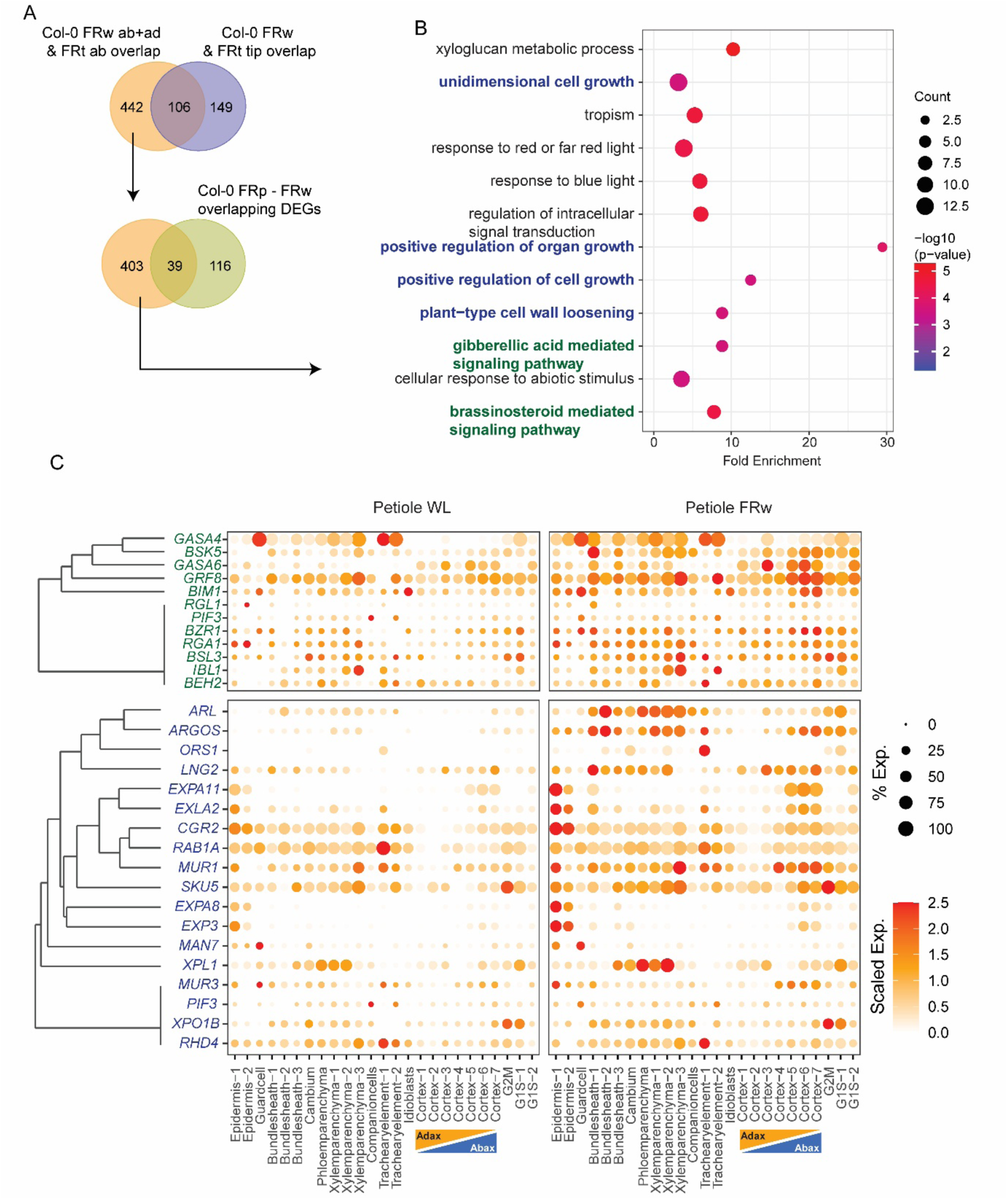
Combinatorial phenotypic responses were associated with additional hormonal and growth regulation transcript changes in the petiole. (A). Venn diagrams of DEGs from both bulk RNAseq experiments to filter DEGs linked to enhanced petiole elongation during FRw enrichment. (B) GO analyses of 403 DEGs resulting from the previous filtering. (C) Average gene expression and percentage of cells expressing the DEGs per cluster in petiole WL and FRw enrichment, stemming from the GO categories highlighted in corresponding green and blue for GA and BR and growth regulation respectively. Gene clustering was based on their cell-type-specific differential expression pattern, where the differential expression was encoded as a binary variable (non DEG = 0, DEG = 1). Genes were then grouped by hierarchical clustering using the binary distance (Jaccard). Colour scale represents average level of expression per gene, set from 0.0 to 2.5 max. Size of the dot represents the percentage of cells in a cluster expressing the gene in question.

## Discussion

### Unidirectional elongation and differential growth are controlled by spatially distinct networks

Hyponasty is induced upon signalling of FR-enrichment in the leaf tip (Pantazopoulou et al., 2017; Michaud et al., 2017; Küpers et al., 2023; Oskam et al., 2024; Supplemental Figure 1A), whereas unidirectional petiole elongation is induced in part by local FR enrichment of the petiole itself (Pantazopoulou et al., 2017); Figure 5B). In plants that are exposed to uniform FR light enrichment, i.e. the standard setup in most experimental studies, both these responses are initiated and co-occur. We aimed to unravel transcriptional networks that could spatially coordinate these differential growth processes in leaf petioles. Several gene sets related to different components of the co-occurring responses were defined from bulk RNAseq studies on dissected leaf tissues and subsequently studied with cell type resolution through scRNA sequencing (Figure 4,5, Supplemental Figure 2,7). We observed that expression of genes associated with general cell elongation, occurring in both petiole elongation and hyponasty, is triggered in both epidermis and abaxial cortex clusters, and in the vasculature cell clusters (Figure 4B, 7). Together, these regulations cover most cell types, and this is also what we observed for the gene set that we specifically associate with cell-autonomous low R:FR-induced unidirectional petiole elongation (Figure 7). In contrast to the broad expression of cell elongation networks, hyponasty-specific differential elongation networks were mostly induced in the abaxial cortex, but not in the epidermis or vascular tissues. We therefore propose that spatially distinct transcriptional programs govern petiole growth, with the hyponasty-associated elongation preferentially localized to the abaxial cortex, and the petiole elongation network broadly distributed across cell types. Together they control petiole differential growth and elongation (Figure 7).

**Figure 7.**
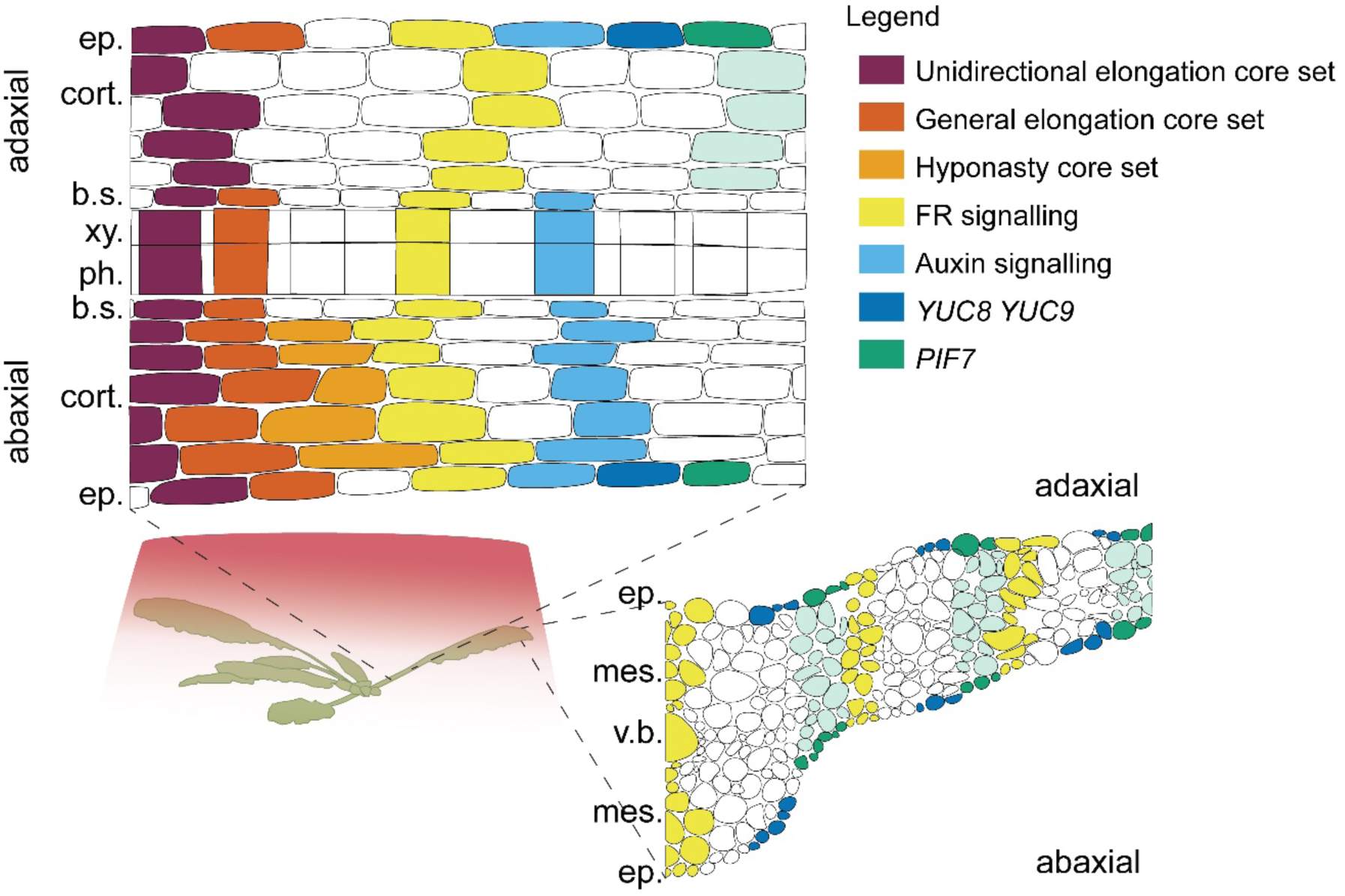
Inferred spatial localization of gene expression during FRw enrichment. Based on the expression of DEGs stemming from the combined bulk RNAseqs and the scRNAseq, we infer the localization of the previously defined core gene sets and genes related to FR signalling, auxin signalling, YUC8 YUC9 and PIF7. Only the cortex (cort.) shows abaxial versus adaxial expression, as for the other tissues, abaxial or adaxial specification could not be inferred from our data. Vasculature is grouped into either xylem (xy.), phloem (ph.) and bundle sheath (b.s.) for the petiole cross section (top left), or as the total vascular bundle (v.b.) in the lamina cross section (bottom right). Epidermis (ep.) is shown without guard cells indicated, and mesophyll (mes.) is not further differentiated. Cell files represent expression throughout the petiole and/ or lamina of: unidirectional elongation core set (purple), general elongation core set (dark orange), hyponasty core set (orange), FR signalling (yellow), Auxin signalling (light blue), YUC 8 and YUC9 (dark blue), and PIF7 (green).

So far, much of our understanding of the regulation of stem elongation is based on observations made in seedling hypocotyls in which tissue-specific interventions in both brassinosteroid and auxin pathways implied that the epidermis is both limiting and regulating the rates of elongation (Savaldi-Goldstein et al. 2007, Procko et al. 2016). This concept is typically extrapolated to other elongating tissues. Indeed, differential elongation between the abaxial and adaxial epidermis has been shown during hyponasty (Polko et al. 2012, Küpers et al. 2023), but this does not proof if the epidermis is driving or following elongation growth. There are examples of leaf bending in other species such as Chinese cabbage (Liu et al. 2024) or rose petals (Cheng et al. 2021a), where cortex cells rather than the epidermis, are proposed to drive the differential growth. Mathematical modelling combined with experimental data suggests the cortex as driving force in male flower anther bending with the epidermis following these growth processes (Silveira et al. 2025). It is possible that also in Arabidopsis, specific regulation in the abaxial cortex of the petiole, on top of cell elongation-facilitative gene regulation across most cell types, determines stronger elongation on the abaxial side as compared to the adaxial side, leading to petiole bending. Thus, with unidirectional petiole elongation-associated gene expression happening throughout the different cell types, we propose that the highly localised expression of hyponasty-associated genes in the abaxial cortex is what might enable differential growth in tissue that is already elongating rapidly under uniform far-red light enrichment.

### Perspectives on the cellular driving forces of differential growth

Cell walls and turgor pressure provide stability and rigidity to plant tissues (Kutschera 1989). Combination of tissue flexibility and differential elongation, cell division, or a combination of the two determines bending potential of an organ (Baskin and Jensen 2013). Mathematical modelling implies that differential elongation of the abaxial petiole alone is sufficient to explain the hyponastic phenotype in ethylene treatments (Polko et al. 2012) that are of roughly comparable magnitude as those induced by light. Because of the combined phenotype of both petiole elongation and hyponasty, FRw enrichment would be expected to induce either substantially more cell elongation compared to supplemental FRt, or additional cell divisions also occur to allow for the average additional 2 mm of elongation in 24 hours of both the adaxial and abaxial petiole sides (Figure 5B). Our results from both the bulk RNA transcriptomic survey and the CYCB1;1-GUS experiment indicated that genes associated with cell division were upregulated in the abaxial petiole during supplemental FRw enrichment (Figure 2C-E). Differential elongation occurs mainly in the basal part of the abaxial petiole during hyponasty (Polko et al. 2012, Küpers et al. 2023). Cell divisions are usually thought to be restricted to the meristematic regions of developing organs (Sablowski and Gutierrez 2022). However, previous research showed *CYCB1* expression until at least day 16 of leaf 8 development throughout the basal region of the petiole (Donnelly et al. 1999). Cell cycle regulation has been implicated before at the adaxial side of the petiole during ethylene induced hyponasty as well (Polko et al. 2015). It may, thus, be possible that cell divisions still occur in the petiole, perhaps especially when cells have become extensively elongated, since cell size is known in meristems to induce cell divisions (Jones et al. 2017). Auxin is known to interact with several regulators of the cell cycle and can promote the progression from Gap 1 (G1) phase to the DNA synthesis-phase (S) (Shimotohno et al. 2021), but its direct influence on cell division under FR enrichment remains open.

### Light signalling and shade avoidance transcriptomes at cellular resolution

We found that both *phyB* and *phyA* photoreceptors in the lamina and petiole are broadly expressed across cell types, including both the outer and inner tissues. Since light quality is unlikely the same across these cell layers, due to especially selective light absorption by chloroplasts in the mesophyll of the lamina, one could ask if photoreceptors in all these tissues can provide information with the same reliability. However, supplemental FR-induced expression of three established R:FR signalling marker genes, *HFR1*, *PAR1* and *ATHB-2*, was observed in all cell types, and comparable between lamina and petiole (Figure 1I, J). This observation indicates that low R:FR signalling, typically through phyB, occurs across most cell types. Furthermore, *PIF4* and *PIF5*, both direct targets of phyB and involved in hypocotyl elongation under light variations and thermomorphogenesis, are also ubiquitously expressed across cell types (Figure 1I, J). However, *PIF7* expression is strongly enriched in the epidermis. PIF7 is the main driver of low R:FR-induced auxin synthesis by inducing expression of genes encoding YUCCA proteins (Li et al. 2012), occurring in the lamina in adult plants (Pantazopoulou et al. 2017). Interestingly, we observed that the PIF7 targets *YUC8* and *YUC9* were entirely restricted to the epidermal cell clusters (Figure 1I, J). Strikingly, the induction of auxin synthesis in the lamina under low R:FR (Michaud et al. 2017, Pantazopoulou et al. 2017, Küpers et al. 2023) (Supplemental Figure 5), was not accompanied by a transcriptional auxin response in this tissue (Supplemental Figure 5, Figure 7). As auxin biosynthesis occurs mostly at the leaf serrations, even in FRw enrichment (Michaud et al. 2017) and is partially transported to the petiole, perhaps only relatively few leaf cells would express an auxin response and this might be difficult to capture. Alternatively, auxin in the lamina might be inactivate, for example through conjugation (Škyvarová et al. 2025) or there could be a mismatch with AUXIN RESPONSE FACTOR (ARF) cellular localisation (Powers et al. 2019).

We conclude that low R:FR signalling occurs across all cell types of the lamina, consistent with phyB and phyA, as well as *PIF4* and *PIF5* expression in all of these layers. However, specification of output, in this example auxin biosynthesis to drive shade avoidance remotely in the petiole, occurs through cell type-restricted expression of the respective set of target regulators, the PIF7-YUC8/9 module. Combining bulk transcriptomics on dissected tissues with scRNAseq thus enabled a novel understanding of physiologically important growth plasticities in response to light signals.

## Materials and Methods

### Plant materials and growth conditions

*Arabidopsis thaliana* plants were grown under WL short day conditions (9 h light, 15 h dark, 70% relative humidity, 20 °C) for 28 days on potting soil. Growing conditions and WL control conditions were 140 µmol m^−2^ s^−1^ Photosynthetically Active Radiation (PAR: 400-700 nm) with a R:FR ratio of 1.5, using Valoya BX-120 modules, NS1 + FR spectrum. Genotypes used in this study were Col-0, *pif4 pif5, yuc8 yuc9, yuc2 yuc5 yuc8 yuc9* (Pantazopoulou et al. 2017), pCYCB1;1::D-BoX:GUS-GFP (Eloy et al. 2011). After 28 days, plants of similar developmental stage were selected for experiments based on the size of the 5th youngest leaf, with a petiole length of 4.5-5.5 mm. For phenotyping experiments, leaves obscuring camera view were removed prior to experimental start time.

### Experimental treatment conditions

Plants were grown 28 days and then exposed to either of the four different light treatments for 24 hours for the phenotypic experiments and 3 hours for the RNA sequencing experiments, starting from ZT 2. These light treatments were a white light (WL) control, and three spatially different far-red light enrichment treatments: Far Red tip (FRt: FR spotlight irradiation of the lamina tip), Far Red petiole (FRp; FR spotlight irradiation of the petiole), and Far Red whole (FRw; exposure of the entire plant to FR enrichment). Local FRt and FRp enrichment were performed using individual L735-06AU LEDs (USHIO, Japan) with a peak emission at 735 nm, resulting in a R:FR ratio of approximately 0.1 at the leaf tip for FRt or the petiole for FRp. Background light was the same as control condition, with a PAR of 140 µmol m^−2^ s^−1^. Supplemental FR for whole plant FR (FRw) was realized using BX-120 LED bars (Valoya) and set to reach a R:FR ratio of 0.1. Background PAR was always maintained at 140 µmol m^−2^ s^−1^.

### Image acquisition and analyses

Leaf angle and elongation measurements were made from images acquired every minute over a 24-hour period by a semi-automated imaging pipeline (Oskam et al. 2024). Coordinates of plant rosette centre, petiole-lamina junction and the leaf tip were determined by visual point selection in the first image. Calculation of angles and length was performed with triangulation between coordinates of the aforementioned points. Data collection was performed in Spyder 5.5.1, with Python 3.9.18. Downstream data analyses were performed in R with R studio, as previously described with updated calculations of the angle, which are deposited on Github (https://github.com/Pierik-Lab) (Oskam et al. 2024). A Tukey test was performed for grouped data of 2-hour blocks to test for statistical differences between comparison groups.

### Bulk RNA sequencing and analyses

For the RNA sequencing transcriptome studies, material of 12 plants was pooled per sample, and three independent biological replicates were collected for further processing. One biological replicate of each treatment x tissue x genotype combination was harvested per day to minimize sampling time. For bulk RNAseq A, petioles were split longitudinally to separate the abaxial and adaxial sides, and the first 5 mm of the leaf was used as leaf tip sample. For bulk RNAseq dataset B, the first two thirds of petioles were harvested whole. Material was snap frozen in liquid nitrogen immediately upon harvesting and stored at -80 °C. Extraction of RNA was done with the RNeasy Mini Kit from Qiagen with on-column DNase treatment. Both mRNA library preparation via polyA-enrichment and sequencing were performed by Novogene (www.novogene.com). Sequencing was done on Illumina NovaSeq 6000 platform via paired-end sequencing with a read length of 150 bases, providing a minimum of 6 GB raw data per sample.

Checking of raw sequence quality was done with FastQC 0.11.8 (http://www.bioinformatics.babraham.ac.uk/projects/fastqc/). Trimming of reads and quality checking was done using TrimGalore 0.6.10 (https://github.com/FelixKrueger/TrimGalore), with default parameters. Paired reads were mapped to the Arabidopsis TAIR10 genome using HISAT2 2.2.1 (Kim et al. 2019), with default settings. The count tables for different feature levels were obtained from the bam files with featureCounts 2.0.6, while considering the Araport11 gene model (Cheng et al. 2017) from Phytozome (https://phytozome-next.jgi.doe.gov/info/Athaliana_Araport11). DEGs were identified using the R package limma according to (Jo et al. 2025a). Counts per million (CPM) were obtained with Bioconductor package edgeR 4.4.2 (Robinson et al. 2010, Lun et al. 2016). Genes were filtered for having at least 1 CPM in at least 3 samples. For every time tissue, we compared the far-red enrichment-treated sample (FRw, FRt or FRp) against the equivalent tissue in WL using edgeR and differentially expressed genes (DEGs) were determined as genes with an Benjamini-Hochberg adjusted P value equal or smaller than 0.01 and a log2FC equal or greater than 0.585 (up regulated genes) or equal or smaller than -0.585 (down regulated genes). Analyses were done in R Studio with R version 4.4.1.

### Single Cell RNA sequencing and analyses

Protoplasts were obtained from leaves and petioles of plants in 3 hours of FRw or WL treatment. The isolation of protoplasts was performed as previously described (Jo et al. 2025b), with modifications. In summary, the lamina and petioles of the 5^th^, 6^th^ and 7^th^ youngest leaves were manually dissected, cut into 1-2 mm slices and vacuum infiltrated for 15 minutes immersed in an enzyme solution (0.4 M D-Mannitol, 20 mM KCl, 20 mM MES, 10 mM CaCl2, 0.1% (w/v) BSA, 1.5% (w/v) Cellulase R10 (Duchefa), 0.5% (w/v) Macerozyme R10 (Duchefa), 5% (v/v) Viscozyme® L (Sigma)). We supplemented the enzyme solution with 25 µM ActinomycinD to block transcription during the protoplast isolation process (Tenorio Berrío et al. 2025). We observed that ActinomycinD was able to supress the activation of some protoplast induced genes without affecting the expression of FR inducing genes (Supplemental Figure 9) (Tenorio Berrío et al. 2025). The samples were kept in the enzyme solution for 2 h under 40 rpm in the dark, and the resulting protoplast solution was filtered through a 40 µm mesh and centrifuged for 5 min at 200 x g at 4°C in a swinging bucket centrifuge with the de-acceleration set to minimal. Protoplasts were resuspended in cold 2 mL of resuspension solution (0.4 M D-Mannitol, 20 mM KCl, 20 mM MES, 1 mM CaCl2, 0.1% (w/v) BSA). The protoplasts were purified through a three-phase density gradient using Optiprep™ (Stemcell™ Technologies) (Jo et al. 2025b). The final protoplast resuspension was counted, and 25,000 cells were loaded on separate lanes of a BD HT Cartridge. Single-cell library preparation was performed using the v3 chemistry of the BD Rhapsody Kit following manufacturer’s instructions. Library concentrations and integrity were accessed in a Bioanalyzer and pooled for sequencing by Useq (Utrecht, Netherlands) on a NovaSeq X flow cell (Illumina).

Raw reads were demultiplexed and their respective single-cell indexes were obtained using bcl2fastq with the following parameters: --use-bases-mask=Y151,I8N9,N8,Y151, --minimum-trimmed-read-length=8 , --mask-short-adapter-reads=8. Processed reads were then aligned to the Arabidopsis genome (TAIR10) and Araport11 gene model (https://phytozome-next.jgi.doe.gov/info/Athaliana_Araport11) using STARsolo with the following parameters: --soloType CB_UMI_Complex, --soloAdapterSequence GTGANNNNNNNNNGACA, --soloCBposition 2_-9_2_-1 2_4_2_12 2_17_2_25, --soloUMIposition 3_10_3_17, --soloCBmatchWLtype 1MM, --soloCellFilter EmptyDrops_CR, --soloUMIlen 8. Filtered count matrices were later analyzed using Seurat version 5.0.2 (Hao et al. 2023).

For each individual sample low-quality cells were removed if mitochondrial or chloroplast gene counts exceeded 5% of total counts. For each tissue (lamina and petiole), FRw and WL samples were integrated using the IntegrateLayers() function from Seurat using the canonical correlation analysis (CCAIntegration) method (Butler et al. 2018). Uniform manifold approximation and projection (UMAP) were calculated using 30 principal component dimensions, and cell-clusters were identified using the Seurat ‘FindClusters’ function with the resolution parameter set to 0.5. Markers for each UMAP cluster were identified using the using the Seurat ‘FindAllMarkers’ and filtered for an average log2FC greater than or equal to 0.25, minimal percent difference of 0.2, and adjusted P values lower than or equal to 0.01. To infer the cell identity of identified clusters, we compared the marker genes to the ones identified in the single-cell transcriptome atlas of the Arabidopsis Leaf (Tenorio Berrío et al. 2025).

For the DEG analysis in the clusters identified in the single cell, we compared the expression of genes between cells of the FRw and WL samples in each respective cell cluster using the ‘FindMarkers’ function from Seurat with the method MAST (Finak et al. 2015)). We consider DEGs every gene with a log2FC equal or greater than 0.25 and an adjusted P value equal or lower of 0.01.

For the classification of adaxial and abaxial identities in the petiole cortex cells, we used the CellCycleScoring function of Seurat and using the top 20 up and down DEGs (based on their significance (adj.P.Val)) found in the bulk RNA-seq when comparing the adaxial versus abaxial samples in WL.

## Data availability

The datasets reported in this paper have been deposited EMBL-EBI BioStudies/ArrayExpress and will be made available upon publication.

## Data visualization: Venn diagrams, GO analyses and heatmaps

Venn diagrams were created using lists of DEGs obtained from the contrast matrices using an online Venn creator: https://bioinformatics.psb.ugent.be/webtools/Venn/. GO enrichment analyses were performed using PANTHER version 19. Figures were created in R Studio with R version 4.2.3, using packages ggplot2, dplyr, tidyr and readxl. Resulting figures were aesthetically improved in Adobe Illustrator 28.6 for fonts and graph size. Clustering was done using Euclidean distance and complete-linkage clustering.

## GUS staining and light microscopy

28 Days old plants pCYCB1;1::D-BoX:GUS-GFP (Eloy et al. 2011) were treated with either WL or FRw starting from ZT2. Whole leaves were harvested at ZT5 and ZT26 and immediately fixated in 90% acetone at -20 °C, and washed twice in washing solution (phospH-Pi pH 7.0, EDTA, K3Fe(CN)6, K4Fe(CN)6.3H2O, followed by a 5 minute vacuum infiltration with staining solution (phospH-Pi pH 7.0, EDTA, K3Fe(CN)6, K4Fe(CN)6.3H2O with 5 mg/μL X-Gluc) and further dark incubation at 37 °C overnight. Continued by a 60 minute immersion in acetic acid : EtOH (3:1) and subsequent washing with 70% EtOH for clearing. Microscopy images were taken with a 10x light AXIO microscope (ZEIS). Samples were scanned as well for visualization at 600 dpi with an EPSON V850 photonegative scanner. Microscopy images of petioles were analysed for the number of spots in ImageJ using Colour Threshold and Analyze Particles. Further downstream analyses were done in R Studio with R version 4.2.3.

## Acknowledgements

We thank Yorrit van de Kaa for plant care and Andrés Romanowski for support with initial RNAseq analyses.

