## Supplemental Figures 1-9 for "Local and distal signals drive cell-type-specific leaf responses during shade avoidance"

Supplemental figures belonging to: Local and distal signals drive cell-type-specific petiole responses during shade avoidance

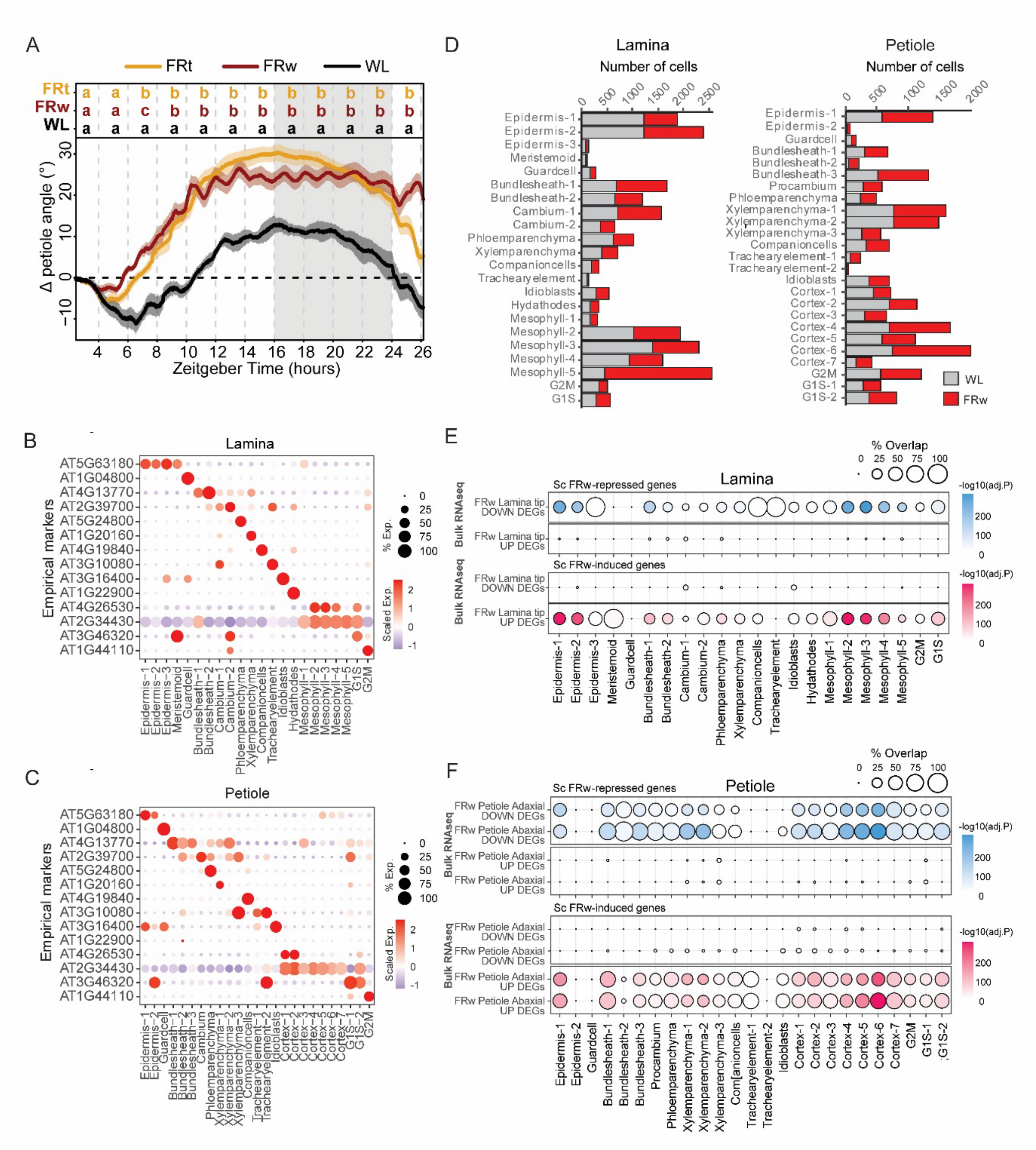


*Supplemental Figure 1. (A) Phenotypic response of FR enrichment on the leaf tip (FRt) and whole rosette FR enrichment, compared to WL control conditions. Petiole angle change over a 24 hour time course, measurements where done every minute, treatment was started at ZT2. WL n=5, FRt n=9, FRw n=11. Grey area indicates night time without light or FR enrichment. Letters on the top represent whether p<0.05, calculated with 2 hour blocks and a post-hoc Tukey test. (B) Average gene expression of marker genes for different cell type clusters in the lamina and (C) the petiole.  Colour scale represents average level of expression per gene, set from -1.0 to 2.5 max. Size of the dot represents the percentage of cells in a cluster expressing the gene in question. (D) Number of cells per cluster for lamina and petioles in WL (grey) and FRw enrichment (red). (E) Up and down regulated DEGs overlapping with corresponding bulk RNAseq tissue per cluster for the lamina and (F) the petiole.*

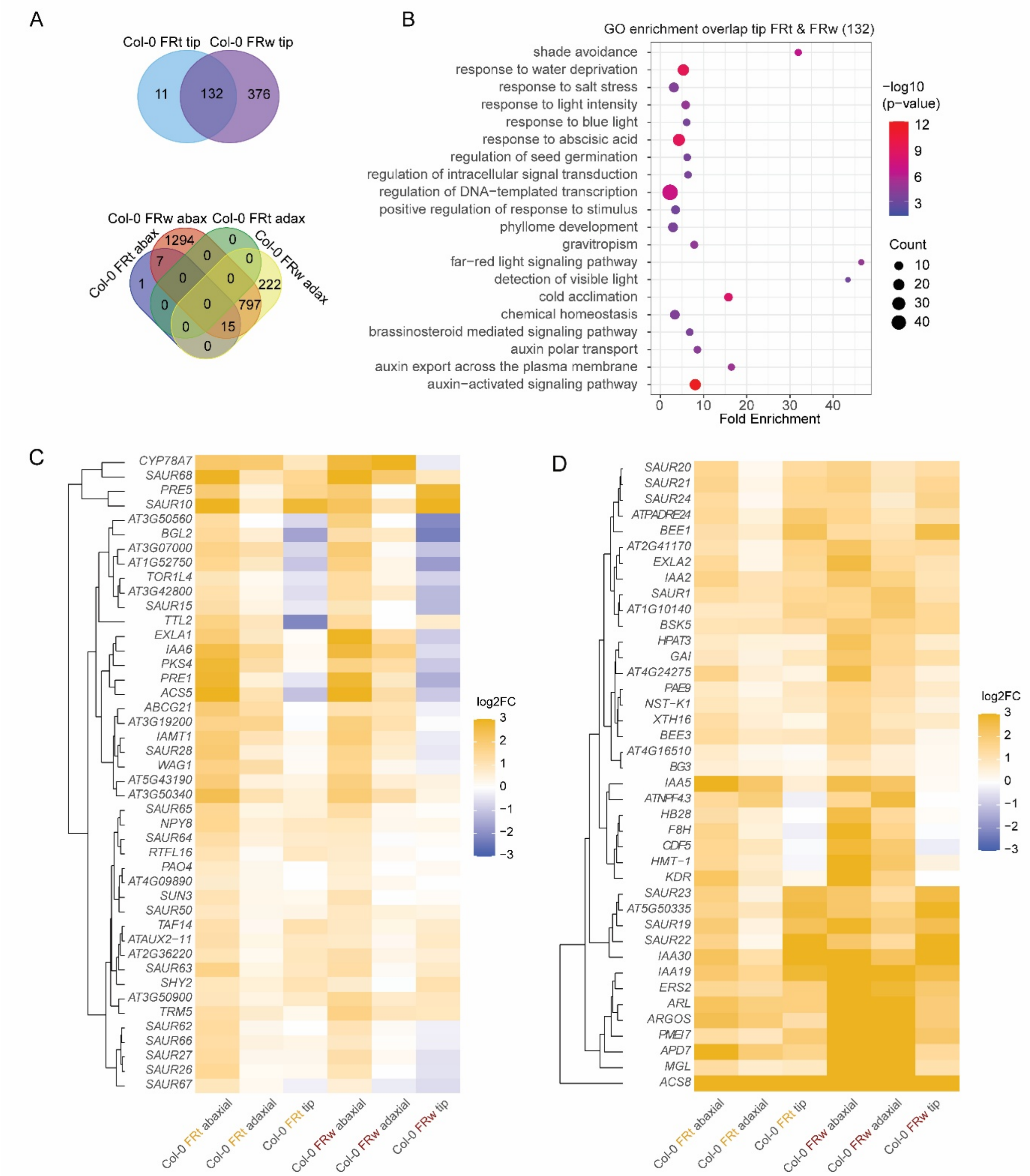


*Supplemental Figure 2. Location of FR enrichment results in a unique response in both gene expression and phenotype. (A) Venn diagram of downregulated DEGs at both the lamina and the petiole. (B) GO enrichment analyses of the shared upregulated genes in the lamina between FRt and FRw enrichment. (C) Change in gene expression of DEGs  the hyponasty core set and (D) the general elongation core set. Clustering based on Euclidian distances. Colour scale represents up and down regulation of genes, and is set to a log2FC of -3 to 3 max.*

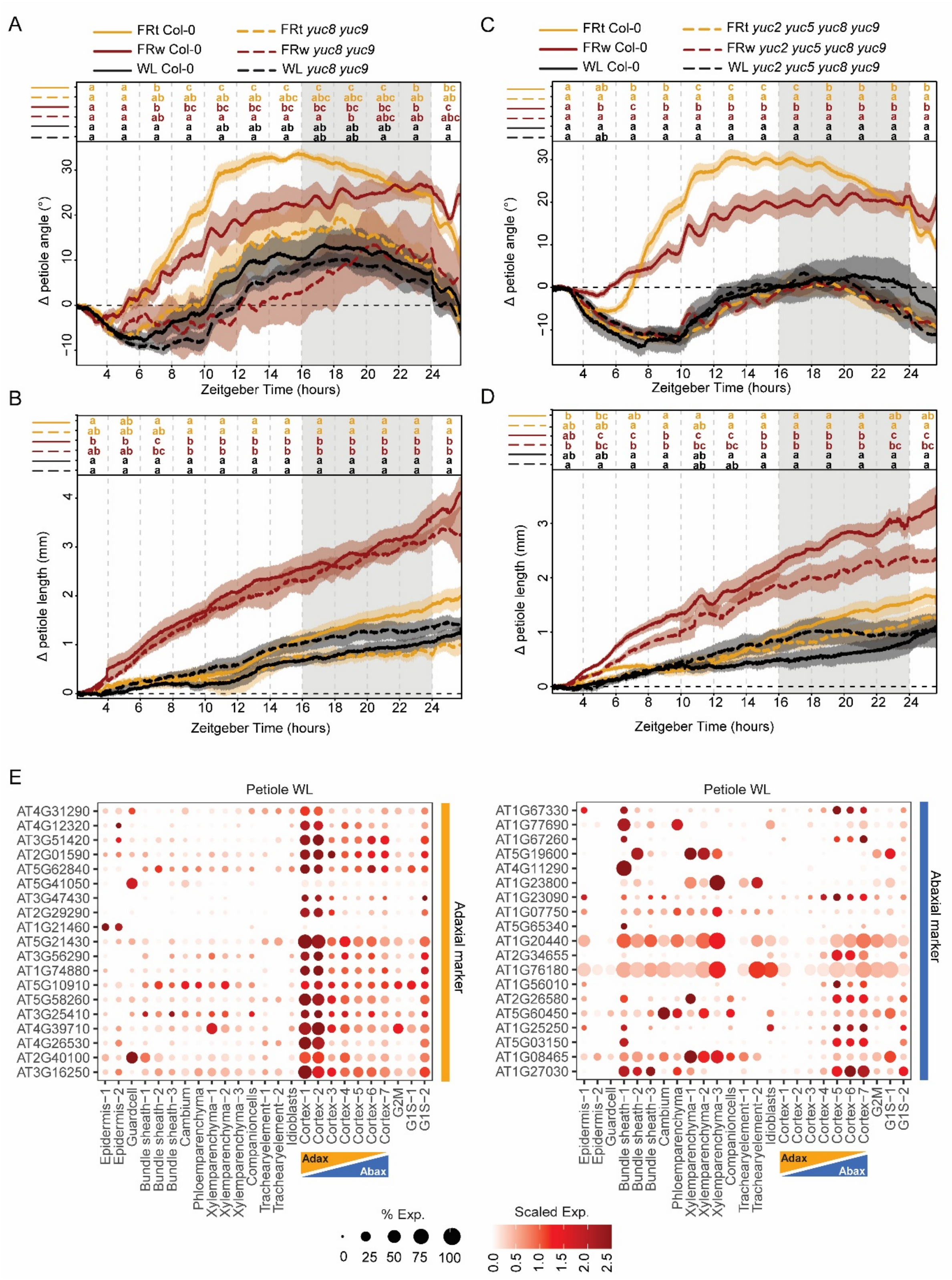


*Supplemental Figure 3. (A) Differential petiole angle, differential leaf angle and petiole elongation are shown for (A,B) yuc8 yuc9 and (C,D) yuc2 yuc5 yuc8 yuc9 in white light (WL), local supplemental FR on the lamina tip (FRt) and supplemental FR on the whole plant (FRw). Plants were measured for 24 hours from onset of treatment. Solid lines indicate Col-0, dotted lines indicate yuc8 yuc9 or yuc2 yuc5 yuc8 yuc9. Replicate numbers as follows: Col-0 in A,B: WL n = 5, FRw n = 7, FRt n = 4; yuc8 yuc9 WL n = 6, FRw n= 5, FRt n = 4. Col-0 in C,D: WL n = 3, FRw n = 11, FRt n = 7; yuc2 yuc5 yuc8 yuc9 WL n = 6, FRw n = 11, FRt n = 7. Grey areas indicate night without light or FR exposure. Letters indicate p < 0.05, calculated per every 2 hours using two-way ANOVA and Tukey post-hoc test. Tissue specific expression pattern of top genes associated with adaxial or abaxial tissue based on the bulk RNAseq. Petiole WL*

*expression in different tissue clusters of top 20 marker genes selected from manually dissected petioles and bulk RNAseq results. Colour scale represents average level of expression per gene, set from 0.0 to 2.5 max. Size of the dot represents the percentage of cells in a cluster expressing the gene in question.*

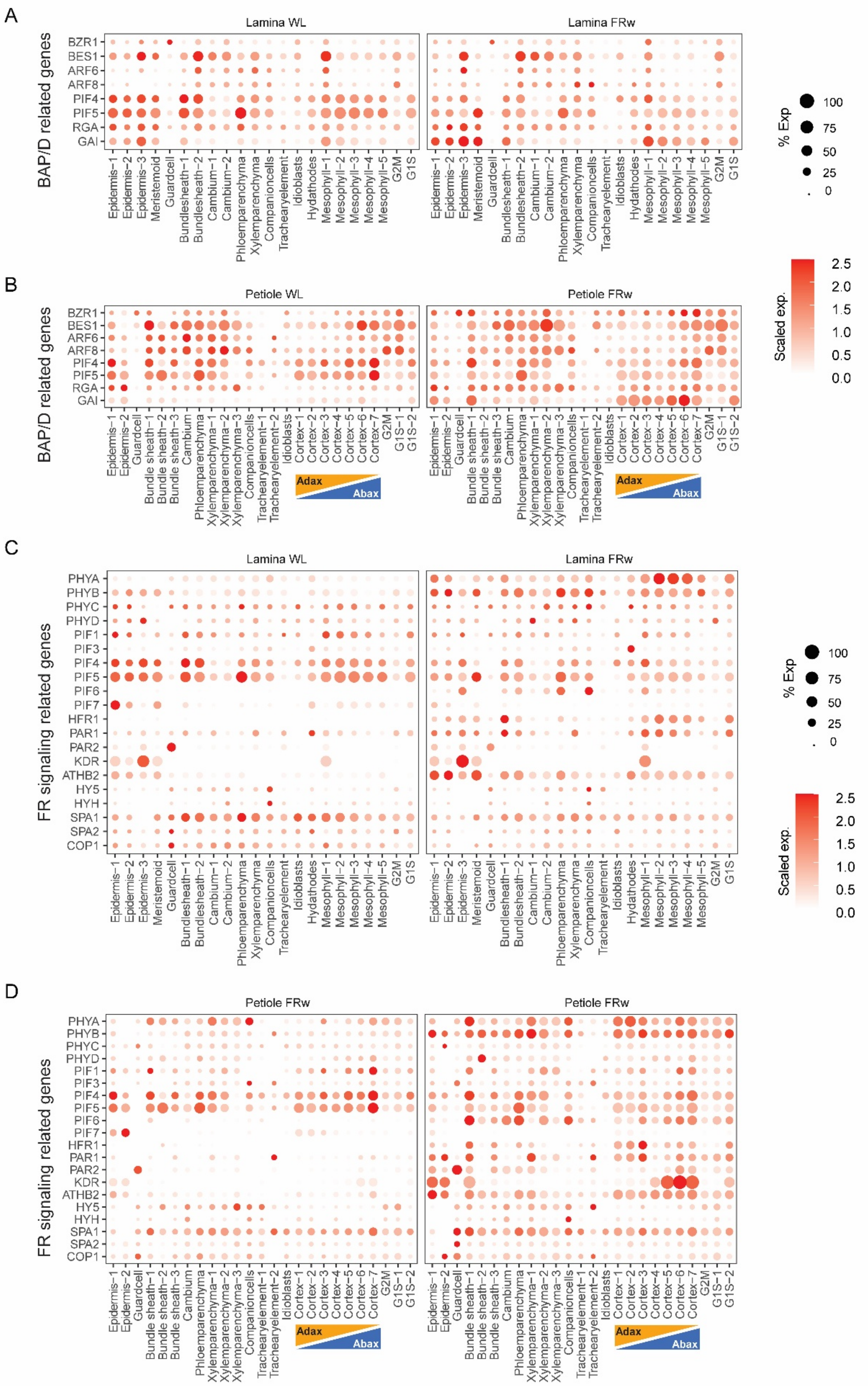


*Supplemental Figure 4. Expression and relative expression within a cluster in WL control and FRw enrichment (A) for genes proposed to be part of the BAP/D module in lamina and (B) the petiole, and (C) genes involved in FR signalling in the lamina and (D) the petiole. Order manually assigned. Colour scale represents average level of expression per*

*gene, set from 0.0 to 2.5 max. Size of the dot represents the percentage of cells in a cluster expressing the gene in question.*

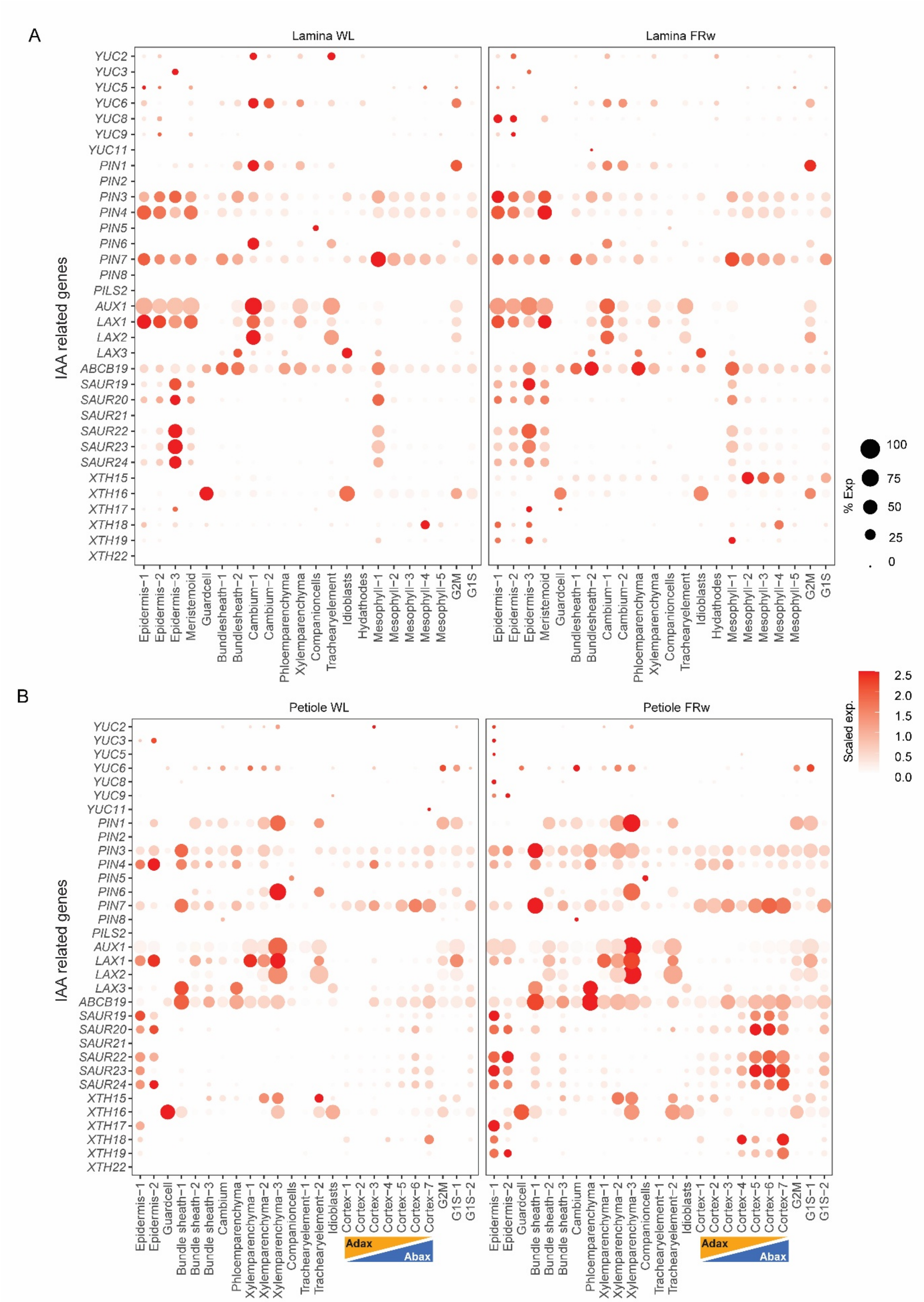


*Supplemental Figure 5. Average expression and relative expression within a cluster of genes related to (A) IAA biosynthesis, transport and regulation of cell expansion though acidification and cell wall modifications in WL and FRw enrichment in the lamina and in (B) the petiole. Colour scale represents average level of expression per gene, set from 0.0 to 2.5 max. Size of the dot represents the percentage of cells in a cluster expressing the gene in question.*

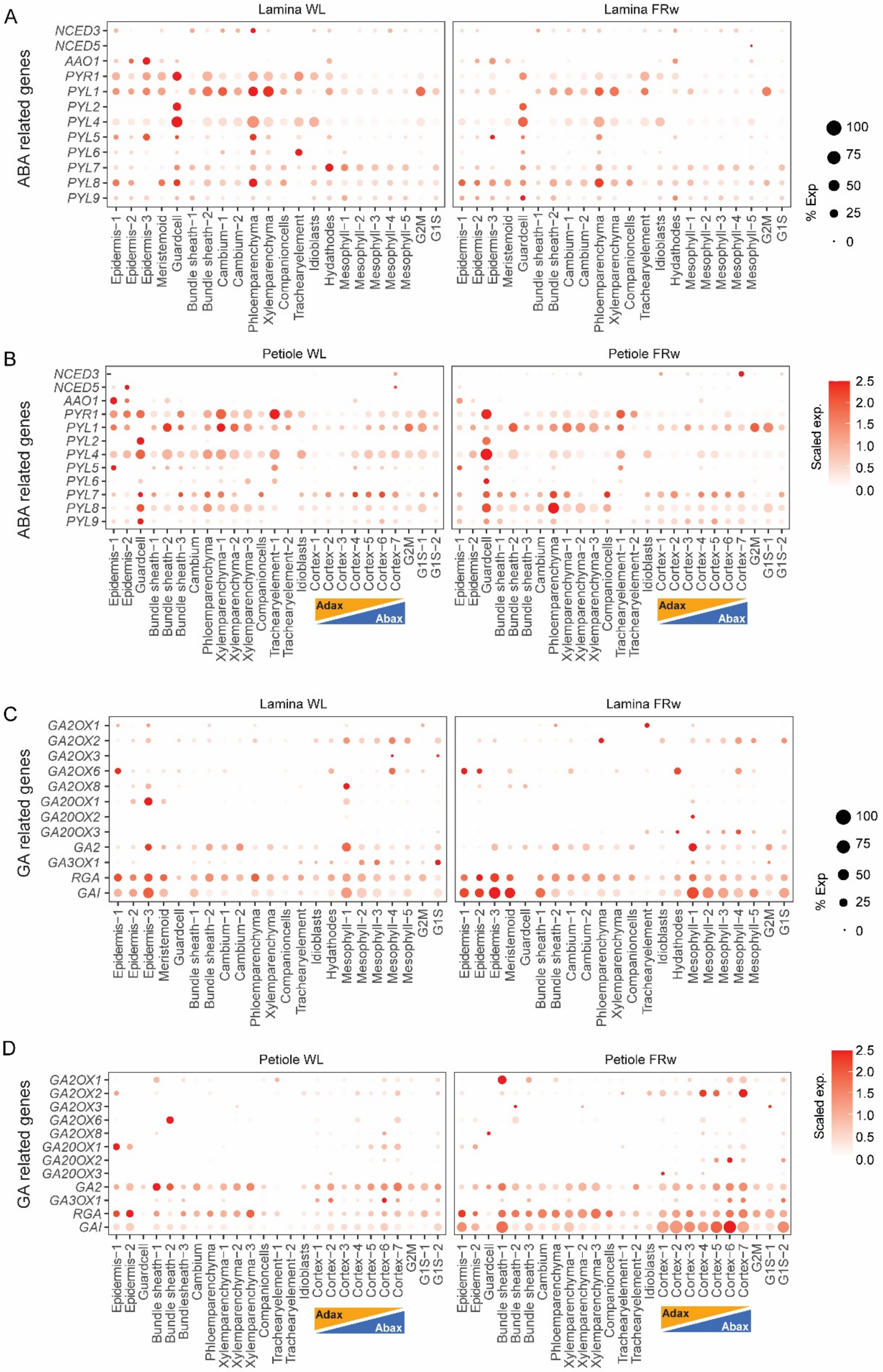


*Supplemental Figure 6. Gene expression and relative expression within a cluster of (A) genes involved in ABA biosynthesis and signalling in the lamina and (B) the petiole. (C) Genes involved in GA biosynthesis and perception in the lamina and (D) the petiole. Both in WL control conditions and FRw enrichment. Order manually assigned. Colour*

*scale represents average level of expression per gene, set from 0.0 to 2.5 max. Size of the dot represents the percentage of cells in a cluster expressing the gene in question.*

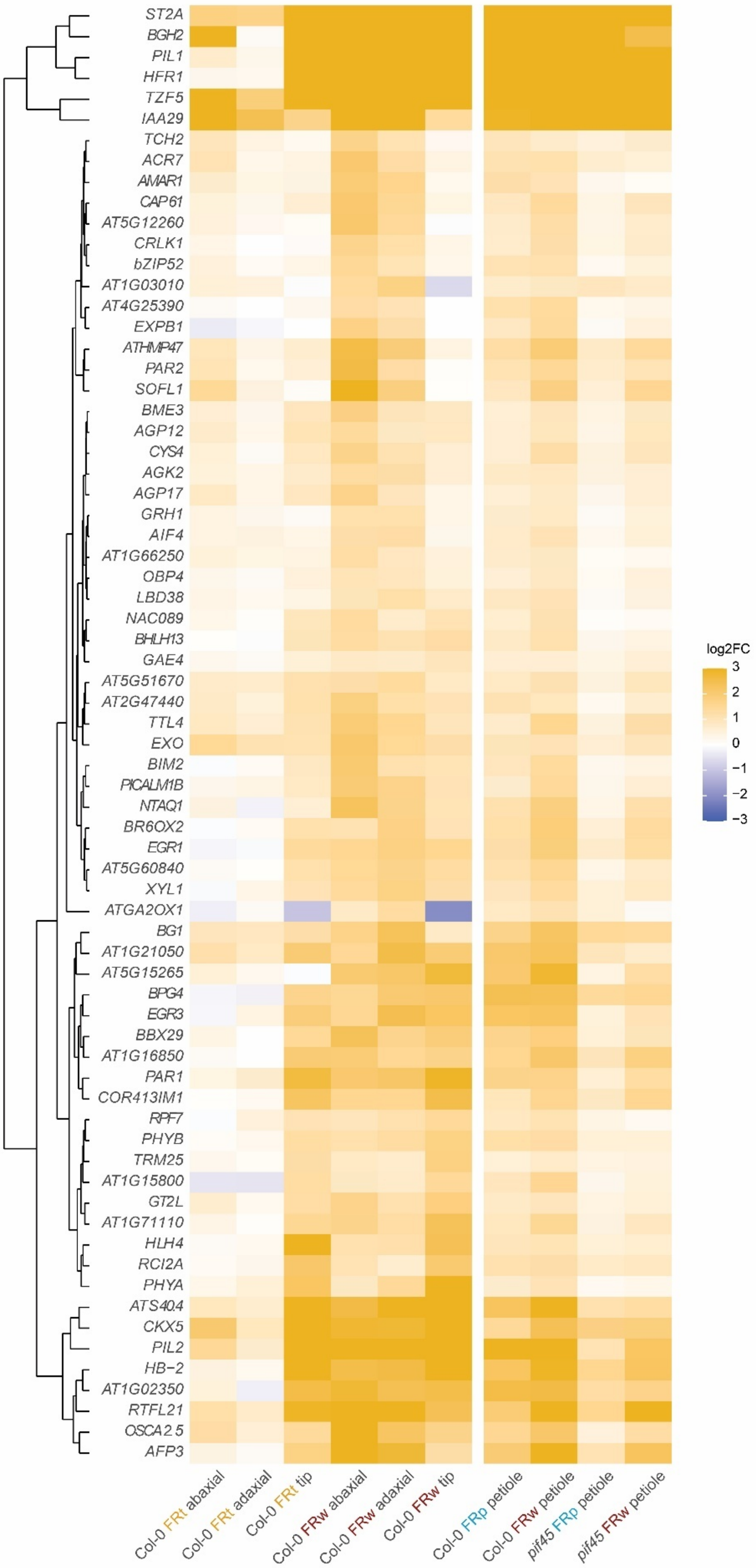


*Supplemental Figure 7. Change in gene expression of DEGs from the elongation core set. The two bulk RNAseqs are separated by a white line. Colour scale represents up and down regulation of genes, and is set to a log2FC of -3 to 3 max.*

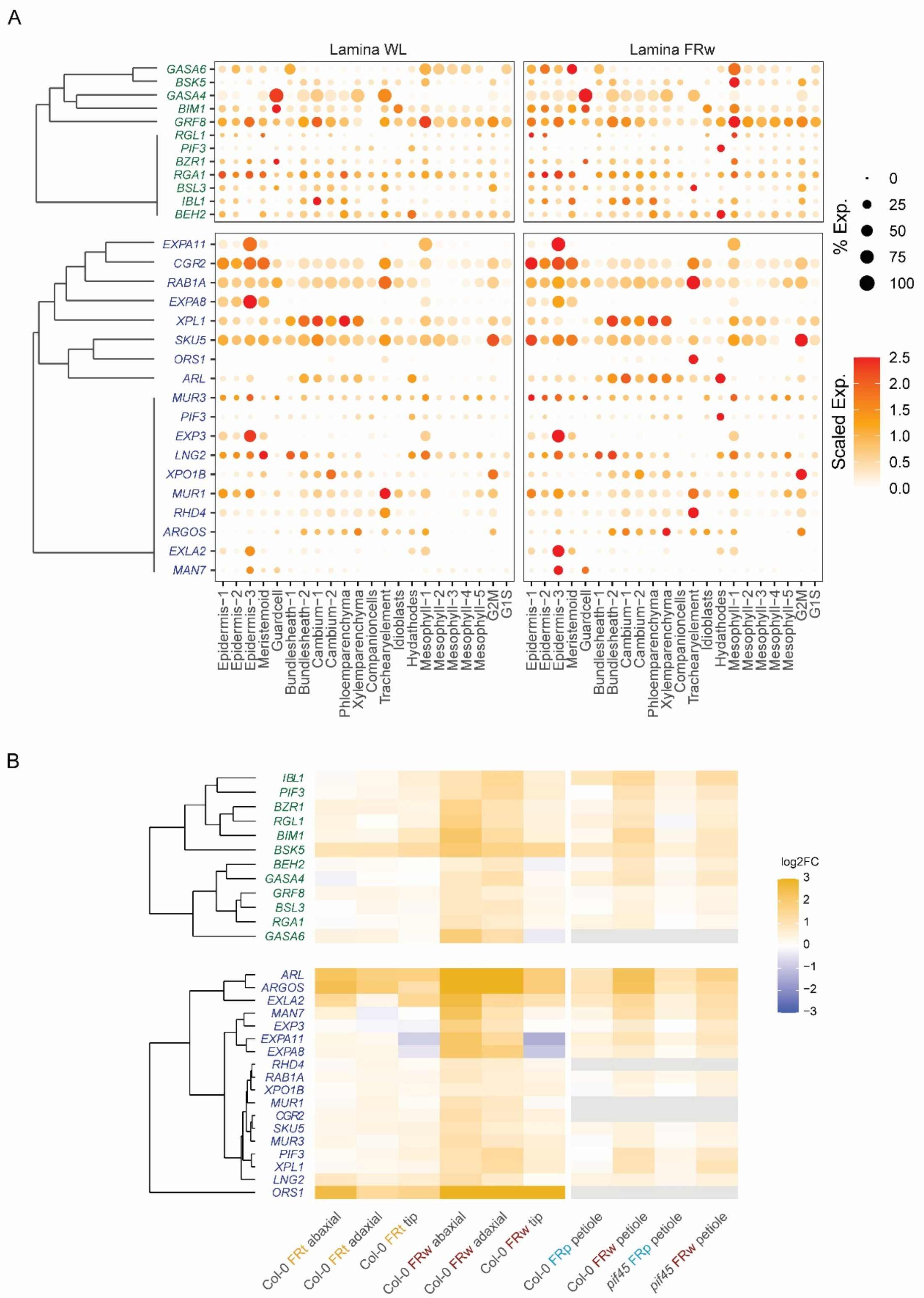


*Supplemental Figure 8. Average gene expression and relative expression within a cluster of genes related to enhanced elongation in the lamina in WL and FRw enrichment per cluster. DEGs stemming from bulk RNAseq filtering of ab/ad DEGs under FRw enrichment. Green represents DEGs related to GA and BR, blue represents genes related to growth regulation. Colour scale represents average level of expression per gene, set from 0.0 to 2.5 max. Size of the dot represents the percentage of cells in a cluster expressing the gene in question. Clustering based on average gene expression, based on Euclidian distances. (B) The same DEGs represented in the bulk RNAseqs. Colour scale represents up and down regulation of genes, and is set to a log2FC of -3 to 3 max, grey indicates lack of gene expression found during this experiment.*

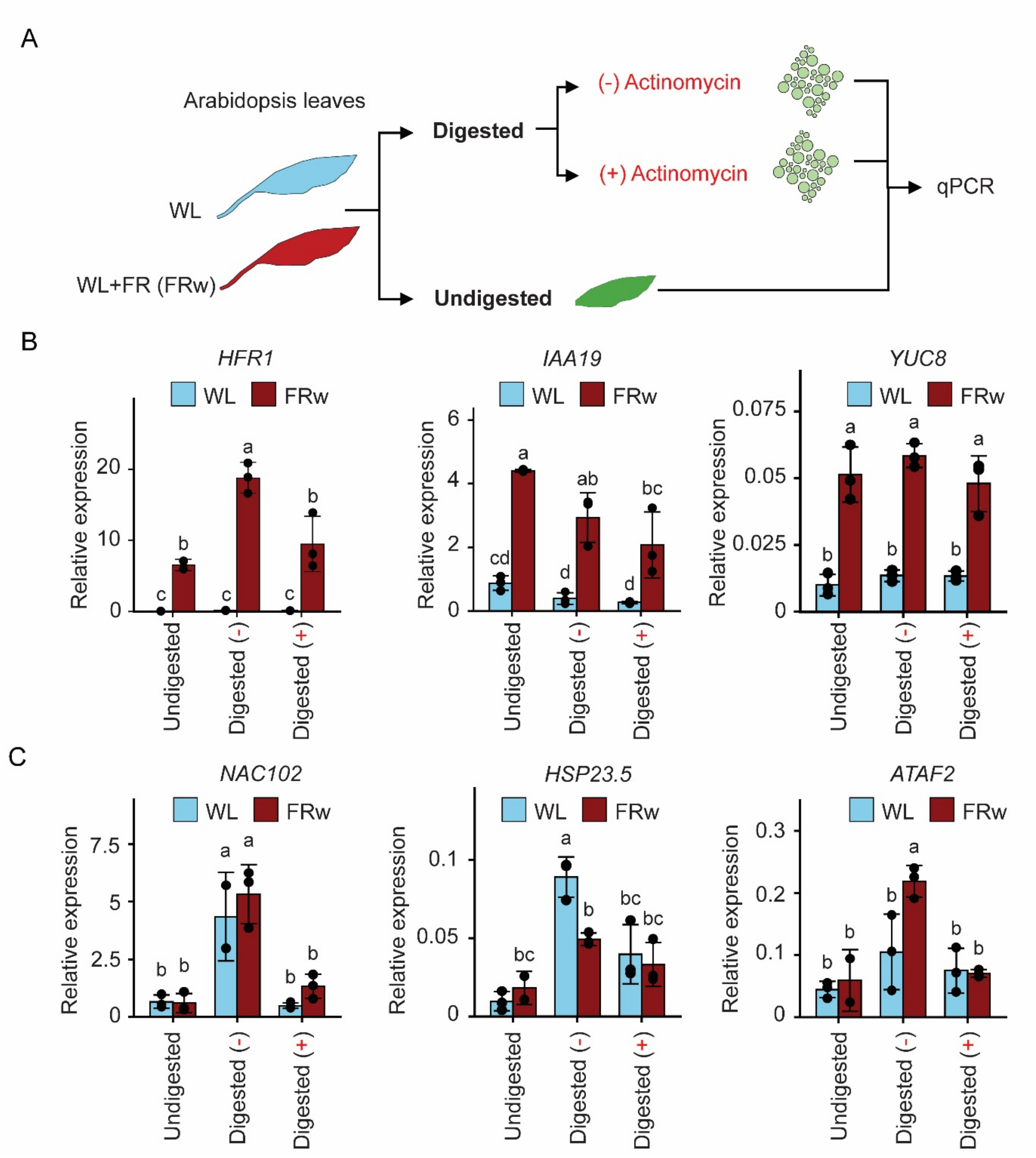


*Supplemental Figure 9. Protoplasting (digestion) and actinomycin did not strongly affect shade-induced transcriptional responses in Arabidopsis leaves. (A) Conceptual diagram of the experiment setting. Arabidopsis leaves from 4-week-old rosettes exposed to WL or FRw conditions (3 hours) were either or not protoplasted (digested) in the presence (+) or absence (-) of actinomycin, followed by gene expression profiling through qPCR. (B) Relative expression levels of FR induced genes in undigested, digested (-) and digested (+) samples. (C) Relative expression of protoplasting induced genes (Tenorio Berrío et al. 2025).*

Tenorio Berrío, R., E. Verhelst, T. Eekhout, C. Grones, L. De Veylder, B. De Rybel and E. Dubois (2025). "Dual and spatially resolved drought responses in the Arabidopsis leaf mesophyll revealed by single-cell transcriptomics." New Phytologist **246**(3).
